# Genomic Characterization of the Endangered Medicinal Polypore Agarikon (*Laricifomes officinalis* syn. *Fomitopsis officinalis*)

**DOI:** 10.64898/2026.06.01.728891

**Authors:** Patrick I. Bennett, Zolton J. Bair, Alexander J. Bradshaw, Paul Stamets

## Abstract

Agarikon (*Laricifomes officinalis* syn. *Fomitopsis officinalis*) is an endangered fungus belonging to a unique lineage in the Polyporales (Basidiomycota) with a growing body of evidence supporting its medicinal value. In this study, we report the hybrid *de novo* assembly and annotation of the first *L. officinalis* nuclear and mitochondrial genome sequences, with a nuclear genome size of 28.76 Mb assembled across 66 scaffolds (51.96% GC content; BUSCO completeness of 99.4%), and a complete core mitochondrial genome size of 197.67 kb. Structural and functional annotation of the nuclear genome yielded 8,717 predicted genes including 8,604 protein-coding genes, with 310 genes in 27 biosynthetic gene clusters. We characterized the mating type loci *matA* and *matB,* consistent with a tetrapolar mating system, and identified genes encoding key enzymes involved in triterpenoid and polyketide biosynthetic pathways that lead to the production of a diverse array of secondary metabolites. Additionally, we conducted maximum likelihood phylogenomic analysis to confirm the taxonomic position of *L. officinalis* among 21 species in Polyporales using protein sequences for 860 shared BUSCO genes. This high-quality annotated genome of *L. officinalis* will serve as a foundation for further investigations into the evolutionary history of this distinct fungal lineage, provide a reference for future population genomic analyses, and elucidate mechanisms underlying the synthesis of the bioactive compounds responsible for agarikon’s wide-ranging medicinal benefits.

## INTRODUCTION

The polypore known colloquially as agarikon, eburiko, and “quinine conk” (*Laricifomes officinalis* (Vill.) Kot. & Pouzar syn. *Fomitopsis officinalis* (Vill.) Bondartsev & Singer) is a flagship species of old growth forest ecosystems and is classified as Endangered on the International Union for Conservation of Nature (IUCN) Red List due to ongoing population decline resulting primarily from overharvesting and habitat loss occurring since the early 20^th^ century (Kałucka and Svetasheva 2019; Girometta et al. 2021; Gilhen-Baker et al. 2022). Agarikon has been recorded across the temperate and boreal forests of the Northern Hemisphere, including North America, Eurasia, and North Africa, with a geographic distribution generally coinciding with those of its coniferous tree hosts including but not limited to larch (*Larix* spp.), Douglas-fir (*Pseudotsuga menziesii*), cedar (*Cedrus* spp.), fir (*Abies* spp.), pine (*Pinus* spp.), spruce (*Picea* spp.), and hemlock (*Tsuga* spp.) (Kałucka and Svetasheva 2019).

*Laricifomes* represents a distinct lineage in the Polyporales (Binder et al. 2013; Ortiz-Santana et al. 2013; Liu et al. 2023; Spirin et al. 2024). Despite its previous taxonomic classification, phylogenetic analyses have shown that it does not form a monophyletic group with members of the genus *Fomitopsis*, and thus is now classified as the only known member of the genus *Laricifomes* in a family with the proposed name of Laricifomitaceae (Ortiz-Santana et al. 2013; M.-L. Han et al. 2016; Liu et al. 2023; Spirin et al. 2024). Two other species, *L. maire* and *L. concavus,* had been classified in the genus previously though these are most often considered members of *Fomitopsis* (Cunningham 1950; Buchanan and Ryvarden 1988; Buchanan and Ryvarden 2000; Spirin et al. 2024). The taxonomic position of *L. officinalis* in its own genus and family reflects its evolutionary divergence, suggesting that it may possess unique genomic and ecological adaptations as well as potentially novel physiological and biochemical properties.

Wood decay fungi serve important roles in forest ecosystem processes such as carbon cycling and tree regeneration (Fukasawa 2021; Niego et al. 2023). Brown-rot fungi such as *L. officinalis* produce lignin-rich residues that contribute to the accumulation of organic matter in soils, increasing water-holding capacity, and creating favorable microsites for seedling regeneration (Gilbertson 1980; Fukasawa 2021). As a heart rot fungus, agarikon also promotes the formation of cavities that serve as habitat for many wildlife species including small mammals and birds (Gilbertson 1980; Jackson and Jackson 2004; Fukasawa 2021). In addition to the many ecosystem services provided, *L. officinalis* also has a long history of use for medicinal and spiritual purposes (Gilbertson 1980; Blanchette et al. 1992).

The first written documentation of its use as medicine dates to the first century CE, when Dioscorides Pedanius wrote about “agaricon” as the “elixir of long life” used to treat a variety of illnesses including indigestion, hernias, dysentery, jaundice, asthma, and tuberculosis (Buller 1914; Gilbertson 1980; Dioscorides 2000). *Laricifomes officinalis* has been highly regarded from a pharmacological perspective and historically was recommended for treatment of “all internal disorders” (Buller 1914; Gilbertson 1980; Dioscorides 2000). Agarikon was also used as medicine for centuries in North America, Europe, North Africa, and Asia to treat a wide range of ailments (Gilbertson 1980; Stamets 2005a; Grienke et al. 2014; Muszyńska et al. 2020; Gafforov et al. 2025).

Recently, a placebo-controlled clinical trial demonstrated that a combination of agarikon and turkey tail (*Trametes versicolor*) mycelia cultivated on brown rice (FoTv) was an effective adjunct to vaccination, both reducing side effects and potentiating more durable antibody generation (Saxe et al., 2026), while a related clinical trial demonstrated FoTv’s value as an antiviral treatment, significantly reducing the number and severity of COVID-19 symptoms (Saxe et al., *in preparation*). Although numerous studies have demonstrated the medicinal value of agarikon, the precise mechanisms of action are understudied, and many of the observed benefits have not been attributed to a single bioactive component.

Basidiomycetes, and polypores in particular, are known to produce a wide array of secondary metabolites with demonstrated bioactivities (Lin et al. 2017; Hassan et al. 2022; Du et al. 2024; Karunarathna et al. 2025; Liu et al. 2025). Biosynthetic genes underlying some of these secondary metabolites have been characterized, including the sesquiterpene synthase (STS) genes encoding enzymes involved in the production of structurally diverse bioactive terpenes in polypore fungi (Chen et al. 2022; Dong et al. 2022; Hassan et al. 2022; Wang et al. 2023), as well as polyketide synthase (PKS) gene clusters encoding enzymes involved in the production of bioactive molecules such as laetiporic acid, orsellinic acid, and their derivatives (Yu et al. 2016; Gressler et al. 2021; Zhang et al. 2024).

*Laricifomes officinalis* is now being viewed in a modern pharmacological context, with burgeoning drug discovery research being undertaken to investigate agarikon’s diverse entourage of bioactive molecules for potential applications in the treatment of cancer, viral and bacterial infections, respiratory ailments, and inflammation (Stamets 2005b; Hwang et al. 2013; Girometta 2019; Fijałkowska et al. 2020; Han et al. 2020; Muszyńska et al. 2020; Gafforov et al. 2025). Secondary metabolites produced by *L. officinalis* include lanostane triterpenoids with demonstrated antimicrobial and anti-inflammatory activities (Wu et al. 2004; Wu et al. 2009; J. Han et al. 2016; Han et al. 2020), chlorinated coumarins with anti-tuberculosis activity (Hwang et al. 2013), as well as various sesquiterpenoids, sterols, indoles, polysaccharides, and phenolic compounds with antibacterial, anti-cancer, anti-inflammatory, and antifungal potential (Fijałkowska et al. 2020; Gafforov et al. 2025).

However, the genes encoding the biosynthetic machinery involved in the production of these compounds have not been previously characterized. Here we present the nuclear and mitochondrial genomes of *L. officinalis,* evaluating genome structure and content with an emphasis on secondary metabolite biosynthesis and synteny with closely related polypore fungi. These reference genomes will serve as a valuable foundational resource for further research into the ecology, evolutionary history, and pharmaceutical potential, and will facilitate efforts to conserve the genetic diversity present within populations of this endangered fungus.

## MATERIALS AND METHODS

### Sample Collection, Isolation, and DNA Extraction

The *L. officinalis* isolate sequenced for this study (“F01”) was originally collected in 2001 from a large conk growing on Douglas-fir near Morton, Washington, USA. Hymenial tissue was sampled using a sterile cork-borer and used to isolate a pure dikaryotic mycelium culture on malt extract agar (MEA) amended with 50 mg/L gentamycin sulfate. Live cultures were maintained on MEA in refrigerated slants and preserved for long term viability in cryogenic storage (Zalesky et al. 2024). For this study, mycelial tissue for DNA extraction was grown in malt extract broth and harvested via filtration. Genomic DNA was extracted from mycelial tissue using the E.Z.N.A. Plant and Fungal DNA kit (Cat. No. D3485-01, Omega Bio-Tek, Norcross, GA, USA) following manufacturer instructions.

### DNA Sequencing

DNA extracts were sent to Novogene Corporation Inc. (Sacramento, CA, USA) for library preparation and whole genome sequencing. To enable a hybrid *de novo* genome assembly, long reads were sequenced on the PromethION platform (Oxford Nanopore Technologies plc, Oxford, UK), while the NovaSeq 6000 platform (Illumina, Inc., San Diego, CA, USA) was used to sequence 150-bp paired-end short reads.

Illumina sequencing libraries were constructed using the NEBNext Ultra II DNA Library Prep Kit (New England Biolabs Inc., Ipswich, MA, USA), including end repair, dA-tailing, and ligation of NEBNext adapters. Fragments were enriched using PCR with P5 and indexed P7 oligos. After libraries were purified, a Qubit fluorometer (Thermo Fisher Scientific, Waltham, MA, USA) was used to determine concentrations and an Agilent 2100 bioanalyzer (Agilent Technologies Inc., Santa Clara, CA, USA) was used to assess insert size. Quantitative real-time PCR was performed to determine effective concentration prior to pooling libraries.

For long-read sequencing, high molecular weight DNA was size-selected, end-polished, nick repaired, A-tailed, and adapters were ligated using the Oxford Nanopore Ligation Sequencing Kit 1D (Oxford Nanopore Technologies plc, Oxford, UK). After purification with AMPure XP beads (Beckman Coulter, Brea, CA, USA), concentration was measured using a Qubit fluorometer and size distribution was assessed with a bioanalyzer, before quantified libraries were pooled for sequencing.

Quality control performed by Novogene on sequence data yielded reads free of low-quality base calls and adapter sequences. Raw read quality was further evaluated with SeqKit v2.12.0 (Shen et al. 2024), Nanoplot v1.41 (De Coster and Rademakers 2023) for ONT reads, and fastqc v0.12.1 (Andrews 2010) for Illumina reads.

### Genome Assembly

Hybrid *de novo* genome assembly was performed using MaSuRCA v4.1.4 (Zimin et al. 2013; Zimin et al. 2017) with both ONT long reads and Illumina short reads. Preliminary assembly quality assessments were performed using QUAST v5.3.0 (Mikheenko et al. 2018) and BUSCO v6.0.0 (Tegenfeldt et al. 2025) prior to segregating mitochondrial components from the nuclear genome.

Two of the MaSuRCA-assembled scaffolds were identified as being of mitochondrial origin based on size, read depth, GC content (%), as well as homology with publicly available fungal mitochondrion genome sequences (assessed via BLAST+ v2.17.0). Preliminary analysis of a putative core mitochondrial scaffold (221 kb) using minimap2 (v2.30-r1287) revealed low ONT read coverage at each end, as well as two duplicated mitochondrial genes, suggesting an erroneous linear assembly. Both mitochondrial scaffolds were excised from the MaSuRCA assembly, and all subsequent analyses handled nuclear and mitochondrial genomes separately.

The core mitochondrial genome was independently reassembled from Illumina reads using GetOrganelle (v1.7.7.1) (Camacho et al. 2009; Langmead and Salzberg 2012; Jin et al. 2020; Prjibelski et al. 2020), with the two MaSuRCA-assembled mitochondrial scaffolds as a reference and the ‘fungus_mt’ preset as seeds. Annotation was performed using MFannot (Lang et al. 2023) with genetic code “4 Mold, Protozoan, and Coelenterate Mitochondrial; Mycoplasma/Spiroplasma” (https://megasun.bch.umontreal.ca/apps/mfannot/). The remaining 32-kb scaffold, putatively of mitochondrial origin, was evaluated independently and annotated with a combination of MFannot and HHpred (Zimmermann et al. 2018) before visualization with OGDRAW (v1.3.1) (Greiner et al. 2019).

Illumina reads mapping to the nuclear scaffolds were isolated and subsequently utilized to polish the MaSuRCA nuclear assembly scaffolds with Pypolca v0.4.0 (Zimin and Salzberg 2020; Bouras et al. 2024). The polished nuclear assembly was cleaned and sorted using Funannotate v1.8.17 (Palmer and Stajich 2020). RepeatModeler v1.0.11 (Smit and Hubley 2008) was used for *de novo* detection of repeats and transposable elements and the resulting repeat library was used to soft-mask with RepeatMasker v4.1.2-pl (Smit et al. 2013), including processing with RepeatClassifier to annotate repeats by referencing the Dfam database (Storer et al. 2021).

TelomereSearch (McGowan 2021) was used to scan the nuclear genome scaffolds for at least 5 copies of conserved fungal telomeric motifs within 1 kb of either end while allowing for flexibility in the nucleotide repeat length.

### Structural Annotation

Structural annotation was performed on the soft-masked assembly using Funannotate, which included *ab initio* gene prediction using BUSCO to identify conserved gene models for training AUGUSTUS (Stanke et al. 2006; Hoff and Stanke 2019), glimmerHMM, and SNAP, in addition to self-training with GeneMark-ES (Lomsadze et al. 2005; Ter-Hovhannisyan et al. 2008). Reference proteomes from *Fomitopsis schrenkii* (UP000015241) (Floudas et al. 2012), *Fomitopsis quercina* (syn. *Daedalea quercina*) (UP000076727) (Nagy et al. 2016), and *Rhodonia placenta* (syn. *Postia placenta*) (UP000194127) (Gaskell et al. 2017), were downloaded from UniProt and provided as hints to assist gene prediction. A predicted protein data set from *Taiwanofungus camphoratus* (syn. *Antrodia cinnamomea*), downloaded from NCBI GenBank (GCA_022598655.1) (Chen et al. 2022), was also included as protein evidence. The quality of structural annotations was assessed using BUSCO in protein mode. A diagrammatic representation of the nuclear genome including scaffold structure, gene density by percentage in 50-kb sliding windows, GC content, and repeat content, was produced using R (v4.5.1; R Core Team, 2025) in RStudio (v2025.9.1.401; Posit Team, 2025) using the circlize package (v0.4.18) (Gu et al. 2014).

### Functional Annotation

Following structural annotation, functional annotation was performed using InterProScan5 v5.77-108.0 (Jones et al. 2014), antiSMASH v8.0.4 (Blin et al. 2025), and Funannotate, along with eggNOG-mapper (EggNog v2.1.13) (Huerta-Cepas et al. 2019; Cantalapiedra et al. 2021). Other databases referenced by Funannotate for functional annotations included dbCAN v12.0 (CAZymes), Gene2Product v1.97, GO v2026-01-23 (gene ontology), MEROPS v12.5 (protease identification), PFAM v38.1 (protein families), and UniProt DB v2026_01 (protein homology). Signal peptides were identified using DeepSig (Savojardo et al. 2018).

### Phylogenomics

MycoTools (v1.0.0) (Konkel and Slot 2023) was used to retrieve reference genome assembly and annotation files from NCBI GenBank for 21 polypore species and to construct predicted proteomes for each genome. This selection of reference genome data was manually curated with a diverse representation of wood decay fungi in the Polyporales (Liu et al. 2023) with publicly available genome assembly and annotation data. Each of the predicted proteomes was subjected to BUSCO analysis in protein mode with the polyporales_odb10 lineage data set. BUSCO_phylogenomics (v20260305) was used to identify BUSCOs shared among these taxa and *L. officinalis* F01 and to construct a concatenated supermatrix alignment of amino acid sequences of the shared genes, along with a partition file. IQ-TREE (v3.0.1) (Chernomor et al. 2016; Hoang et al. 2018; Wong et al. 2025 Apr 7) was used to reconstruct a maximum-likelihood phylogeny using the supermatrix alignment and ModelFinder (Kalyaanamoorthy et al. 2017) was used to determine the best-fit substitution model for each partition. Partitions sharing the same substitution model were merged. Branch support was calculated from 1000 ultrafast bootstrap replicates. The resulting maximum-likelihood tree was rooted on *Heterobasidion irregulare* TC 32-1 (NCBI Accession No. GCF_000320585) (Olson et al. 2012) as the outgroup and displayed using ggtree (v3.16.3) (Yu et al. 2017; Xu et al. 2022) in RStudio.

### Synteny Analysis

A synteny analysis was performed using SYNY (v1.3.2) (Li 2018; Buchfink et al. 2021; Kille et al. 2023; Julian and Pombert 2024) to compare genome content, structure, and organization between the *L. officinalis* genome and a publicly available annotated reference genome for *Laetiporus sulphureus* 93-53 (NCBI GenBank, GCF_001632365.1) (Nagy et al. 2016) using all scaffolds larger than 500 kb from both genomes. To identify mating type loci in the *L. officinalis* F01 genome, the annotation file was queried for product names corresponding to genes such as mitochondrial intermediate peptidase (*mip*) for *matA*, or *STE3* for the *matB* locus (James et al. 2013; Dong et al. 2022). Synteny among homologous genes in the mating-type gene clusters was evaluated using clinker (v0.0.32) (Gilchrist and Chooi 2021) via the online CompArative GEne Cluster Analysis Toolbox (CAGECAT, v1.0) (van den Belt et al. 2023). To facilitate synteny analysis of biosynthetic gene clusters (BGCs) among related polypores, genomes of *Lae. sulphureus* 93-53 (GCF_001632365.1) and *T. camphoratus* V5 (GCA_022598655.1) were also evaluated using antiSMASH (v8.0.4) (Blin et al. 2025). The *T. camphoratus* V5 genome annotations (GFF) retrieved from NCBI GenBank included multiple coding sequences associated with the same locus, requiring AGAT (v1.6.1) (Dainat et al. 2026) to identify overlapping genes and retain only the longest isoform prior to synteny analysis with clinker. Target BGC regions were isolated and similarly submitted via CAGECAT for clinker analysis to produce plots to evaluate synteny/collinearity.

## RESULTS AND DISCUSSION

### Sequence Data Quality

Illumina sequencing produced a total of 23,292,072 reads comprising approximately 3.5 Gb, resulting in an average coverage depth of 121.5x with 91.3% of bases exceeding a Q30 Phred score. After filtering to a minimum average read quality threshold of Q7, Oxford Nanopore data contained 115,538 reads (76.4% of total reads generated) with an average length of 4.8 kb, N50 read length of 7,726 bp, and a total of approximately 555 Mb, resulting in an average depth of approximately 19.3x (Supplementary Tables 1 and 2).

### Assembly of the Nuclear and Mitochondrial Genomes

The initial MaSuRCA assembly yielded 68 scaffolds with total length of 29 Mb and N50 of 997.5 kb with the largest scaffold spanning more than 2.8 Mb. Although the average GC content of the total assembly was 51.74%, an examination of per-scaffold GC and Illumina mapped depth revealed the presence of two mitochondrial scaffolds that did not conform to the expected characteristics of nuclear genome content. The remaining 66 scaffolds representing the nuclear genome had an average GC content of 51.96%, average Illumina mapped depth of approximately 79x, and average ONT mapped depth of approximately 14.9x, for a total genome sequencing coverage of approximately 94x.

The core mitochondrial genome reconstructed by GetOrganelle was circular, 197,672 bp in length, with 26.4% GC content. The MaSuRCA assembly yielded a 32-kb linear scaffold (28% GC), putatively of mitochondrial origin, containing 6 tandem copies of nearly identical nucleotide sequence. Although reminiscent of a mitochondrial plasmid, fluctuations in ONT read depth along with multiple sequence alignment of the subunits suggest the 32-kb scaffold may contain assembly artifacts arising from complex nested repeats; therefore, the tandem copy number should be interpreted with caution. Evaluations with MFannot and HHpred verified the presence of an organellar DNA polymerase type B (DNA_pol_B_2; PF03175.19), with detected sequence features including coiled coil and transmembrane segments. While this scaffold likely represents a mitochondrial plasmid, its true structure was not fully resolved.

### Assembly Quality

Downstream analyses treated the nuclear and mitochondrial genome assemblies separately. The nuclear assembly contained 66 scaffolds, and a total length of 28.8 Mb with the largest scaffold being 2.84 Mb. The N50 was 997,481 bp and overall GC content was 51.96% (Table 1, Supplementary Table 3). BUSCO analysis of the nuclear assembly with the basidiomycota_odb10 lineage dataset (N = 1,764 total BUSCO groups searched) indicated 99.4% completeness, 98.6% complete and single copy, 0.8% complete and duplicated, 0.6% missing, and no fragmented BUSCOs (0.0%). Similar analysis with the polyporales_odb10 lineage dataset (N = 4,464) indicated 98.9% completeness, 98% complete and single copy, 0.9% complete and duplicated, 1.1% missing, and 2 fragmented BUSCOs (0.0%). Pypolca polishing corrected 71 substitution errors and two indel errors. All subsequent analyses were performed using the polished genome assembly file.

**Table 1.** Assembly metrics for the *Laricifomes officinalis* F01 nuclear genome, with BUSCO completeness assessed using the basidiomycota_odb10 lineage dataset.

| Metric | Value |
| --- | --- |
| Genome size (bp) | 28,757,392 |
| Scaffolds | 66 |
| N50 (bp) | 997,481 |
| Largest scaffold (bp) | 2,838,896 |
| GC content | 51.96% |
| Complete BUSCO | 99.4% |
| single-copy | 98.6% |
| duplicated | 0.8% |
| Fragmented BUSCO | 0.0% |
| Missing BUSCO | 0.6% |

**Table 2.** Summary of nuclear genome annotations for *Laricifomes officinalis* F01.

|  |  |
| --- | --- |
| **Total genes** | **8,717** |
| Number of predicted protein-coding genes | 8,604 |
| tRNA | 113 |
| **Total Carbohydrate active enzymes (CAZymes)** | **223** |
| Auxiliary activities (AA) | 39 |
| Carbohydrate-binding modules (CBM) | 2 |
| Carbohydrate esterases (CE) | 12 |
| Glycoside hydrolases (GH) | 115 |
| Glycosyl transferases (GT) | 52 |
| Polysaccharide lyases (PL) | 3 |
| **Secondary metabolite (SM) gene clusters** | **27** |
| **Total genes in SM clusters** | **310** |
| **antiSMASH regions** | **23** |
| NRPS-like<sup>1</sup> | 4 |
| Type 1 PKS<sup>2</sup> | 1 |
| NRPS-like, T1PKS | 1 |
| Terpene | 14 |
| Terpene-precursor | 3 |

<sup>1</sup>NRPS = nonribosomal peptide synthase; <sup>2</sup>PKS = polyketide synthase

Two scaffolds in the nuclear genome assembly contained telomeric repeats within 1 kb of both ends, while 23 contained telomeric repeats near one end of the scaffold. Although most assembled scaffolds do not represent full chromosomes, many contain at least one telomere, which suggests a highly contiguous assembly.

### Structural Annotation of the Nuclear Genome

Following RepeatModeler and RepeatMasker, 10.32% of the genome was soft-masked, with 7.41% consisting of long terminal repeat (LTR) elements (Supplementary Table 4). Four small RNAs were identified, accounting for 0.24% of the elements masked by RepeatMasker. Nearly 2% of the elements masked were listed as unclassified, suggesting the presence of species-specific transposable elements that did not match any known repeat families in the Dfam database. Across the 40 largest scaffolds of the genome, regions with the highest density of repeats generally correspond to areas of lower GC content and gene density (Fig. 1a).

**Fig. 1.**
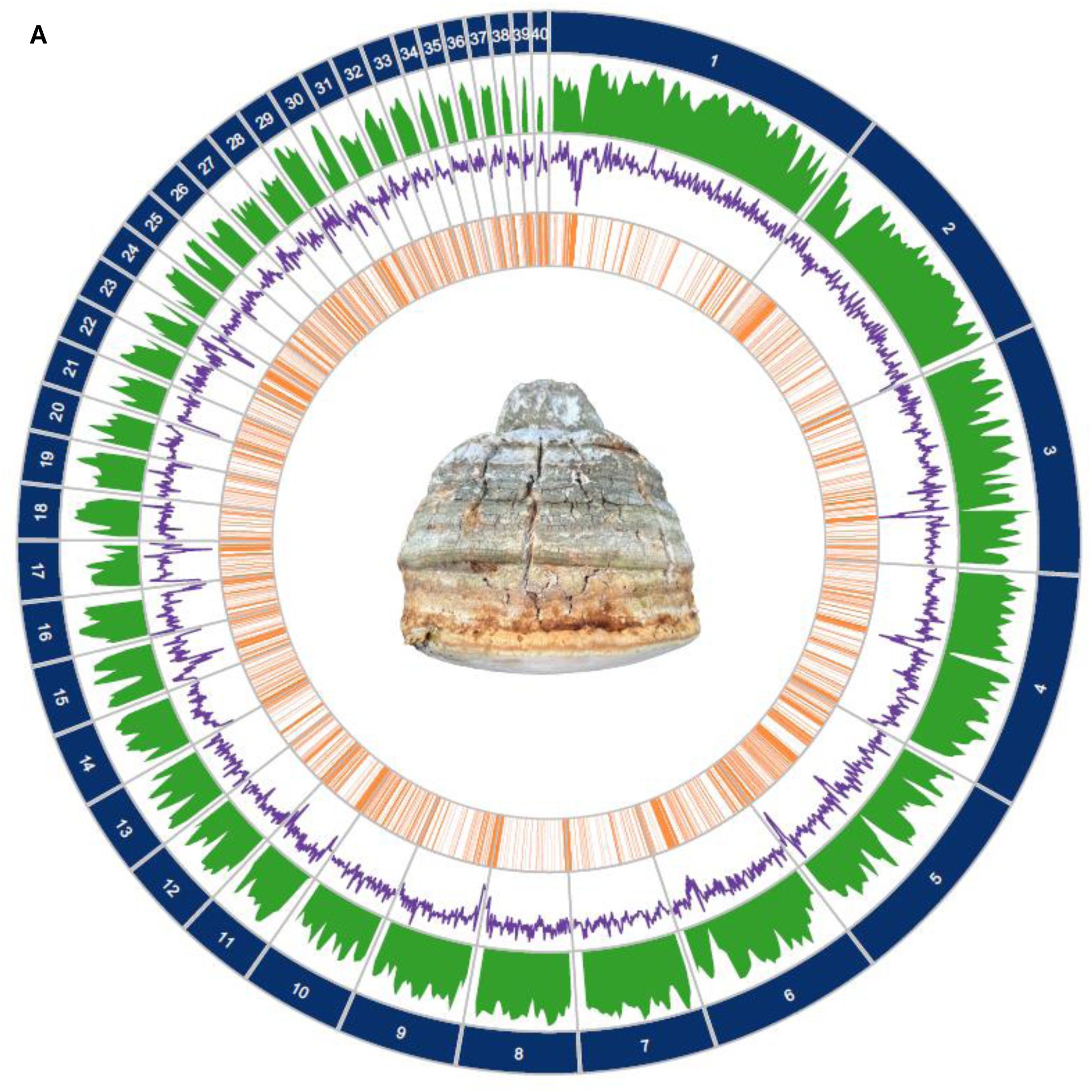
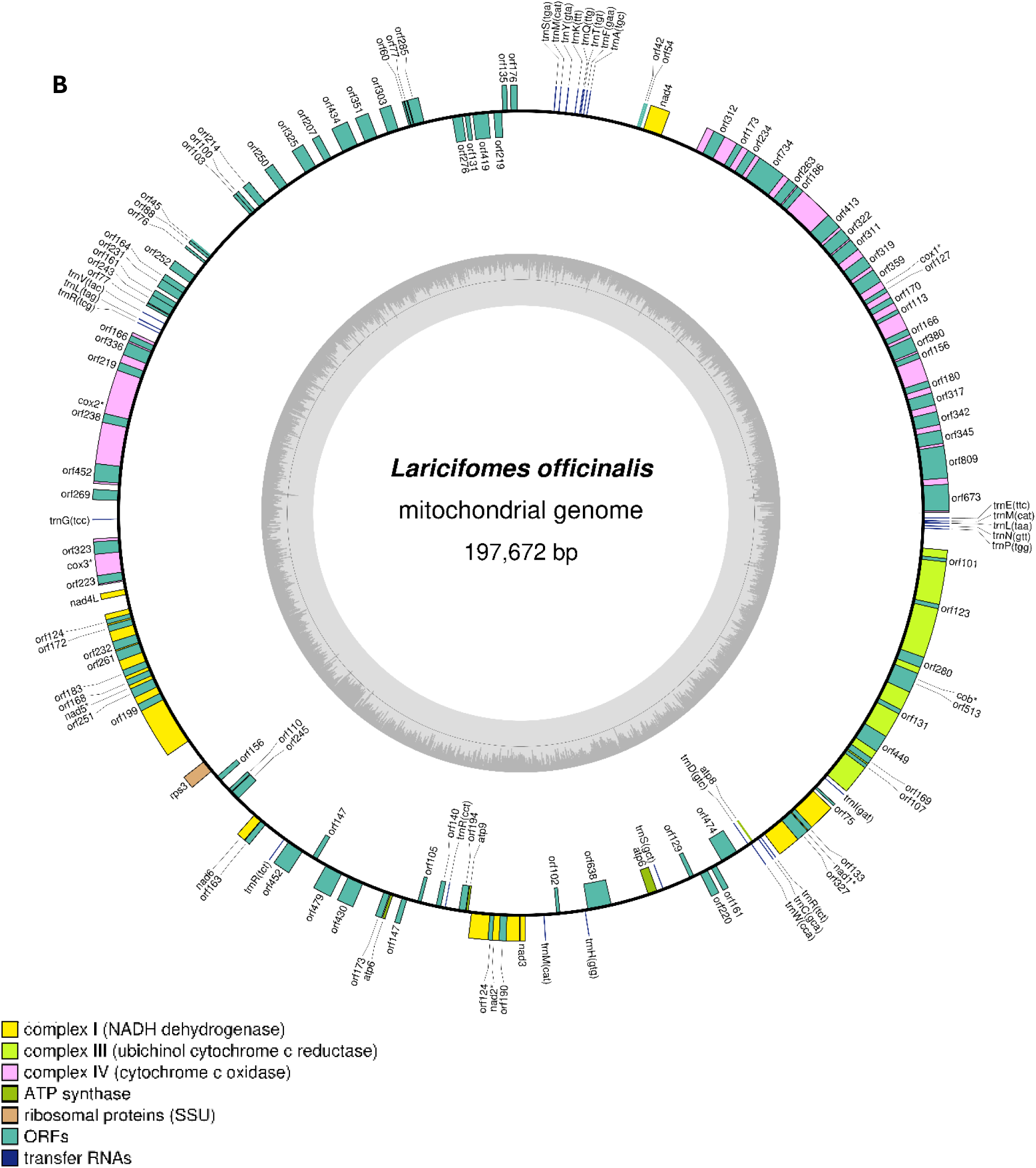
Diagrammatic representations of the *Laricifomes officinalis* F01 genome. a) Nuclear genome. Tracks represent scaffolds (blue), gene density in 50kb sliding windows (green), GC percentage (purple), and repeats (orange). b) Mitochondrial genome. Asterisks indicate genes that contain introns.

Structural annotation revealed that two nuclear scaffolds did not contain any gene content. These scaffolds consisted primarily of repeat content that was masked by RepeatMasker—one was approximately 33 kb with 80.8% repeat content (scaffold_58), while the other was approximately 10 kb with 88.0% repeat content (scaffold_65).

Structural annotation yielded 8,717 total gene models with 8,604 protein-coding genes and 113 tRNAs (Table 2). The BUSCO results for the predicted proteins were C: 93.7% (S: 93.0%, D: 0.7%), F: 3.3%, M: 3.1% for the basidiomycota_odb10 database and C: 90.5% (S: 89.8%, D: 0.8%), F: 3.1%, M: 6.4% for the polyporales_odb10 database.

### Annotation of the Core Mitogenome

Examination of the 197,672 bp circular assembly by GetOrganelle provided strong evidence that this represents the fully resolved core mitochondrial genome. Mitochondrion-specific annotation revealed the core protein-coding genes expected for a fungal mitogenome interspersed with a total of 54 introns, including 2 copies of *atp6*, as well as *atp8*, *atp9*, *cob* (11 introns), *cox1* (23 introns), *cox2* (6 introns), *cox3* (2 introns), *nad1* (4 introns), *nad2* (1 intron), *nad3*, *nad4*, *nad4L*, *nad5* (7 introns), *nad6*, and *rps3*. While mitochondrial rRNA genes (*rns* and *rnl*) were detected, they were not fully annotated by MFAnnot due to ambiguous start and stop boundaries. Partial DNA polymerase genes (*dpo*) were also identified by MFAnnot but were annotated as likely pseudogenes introduced by plasmid integration. Furthermore, there were 24 tRNA genes and 91 unique open reading frames (ORFs) identified, with many representing homing endonuclease genes (HEGs) of the LAGLIDADG and GIY-YIG families (Supplementary Table 5). Numerous ORFs were associated with introns in coding genes, resulting in fragmentation of the gene sequences (Fig. 1b).

The proliferation of introns, in some cases mediated by the actions of HEGs, has contributed to significant expansion of the *L. officinalis* mitochondrial genome. Considerable intron loss and gain events have occurred in the evolutionary history of Polyporales, resulting in highly variable mitogenome sizes, with an examination of 12 species revealing a maximum of 36 introns in the 156 kb mitogenome of *Phlebia radiata* (Salavirta et al. 2014; Wang et al. 2020). At 197.7 kb, the *L. officinalis* mitogenome far exceeds the median for fungi (58 kb) and is currently among the largest mitochondrial genomes reported for basidiomycetes (Ahrendt et al. 2026).

### Functional Annotation of the Nuclear Genome

Functional annotation of the nuclear genome resulted in the identification of 7,958 genes with eggNOG IDs, 5,500 genes with GO terms, 6,736 genes with InterPro domains, 2,501 specific gene products, 440 genes with signal peptides, and 223 genes encoding carbohydrate-active enzymes (CAZymes). Protein-coding genes were functionally annotated with terms corresponding to the Clusters of Orthologous Groups (COG) database, with 6,496 being assigned at least one COG term, including 1,821 with unknown functions (Supplementary Fig. 1). A total of 5,667 genes received at least one Pfam annotation, with a distribution of Pfam domains suggesting investments in signal transduction and stress response via protein kinases (PF00069, PF07714) and fungal-specific transcription factors (PF04082), as well as numerous MFS transporters (PF07690), including some embedded in BGCs which may be involved in the export of bioactive secondary metabolites (Supplementary Fig. 2). Furthermore, 3,640 unique GO terms were assigned across 5,500 genes (Supplementary Fig. 3). A total of 310 genes in 27 biosynthetic gene clusters (BGCs) were identified in 23 regions, including 29 biosynthetic enzymes and 79 smCOGs (Table 2).

### Phylogenomics and Comparative Genomics

A total of 860 BUSCO genes (polyporales_odb10) were shared among all 22 taxa included in the phylogenomic analysis (Suppl. Table 6) and were subsequently incorporated into a concatenated supermatrix alignment of amino acid sequences, containing 468,067 sites, 207,368 of which were parsimony informative. Following ModelFinder, the partitions corresponding to these 860 genes were merged into 45 partitions for further analysis. The maximum-likelihood phylogenetic tree depicted a clade containing *Fomitopsis* species that were formerly included in genera such as *Daedalea*, *Antrodia*, *Piptoporus*, and *Rhodofomes* (Ortiz-Santana et al. 2013; Liu et al. 2023; Spirin et al. 2024), along with a subtending clade containing *Lae. sulphureus* and *L. officinalis* featuring strong statistical support (Fig. 2). Although these species may appear as sister taxa in our phylogenomic analysis, this taxonomic relationship should be interpreted cautiously due to the absence of potentially intermediate taxa that may be more closely related to *L. officinalis* (Liu et al. 2023).

**Fig. 2.**
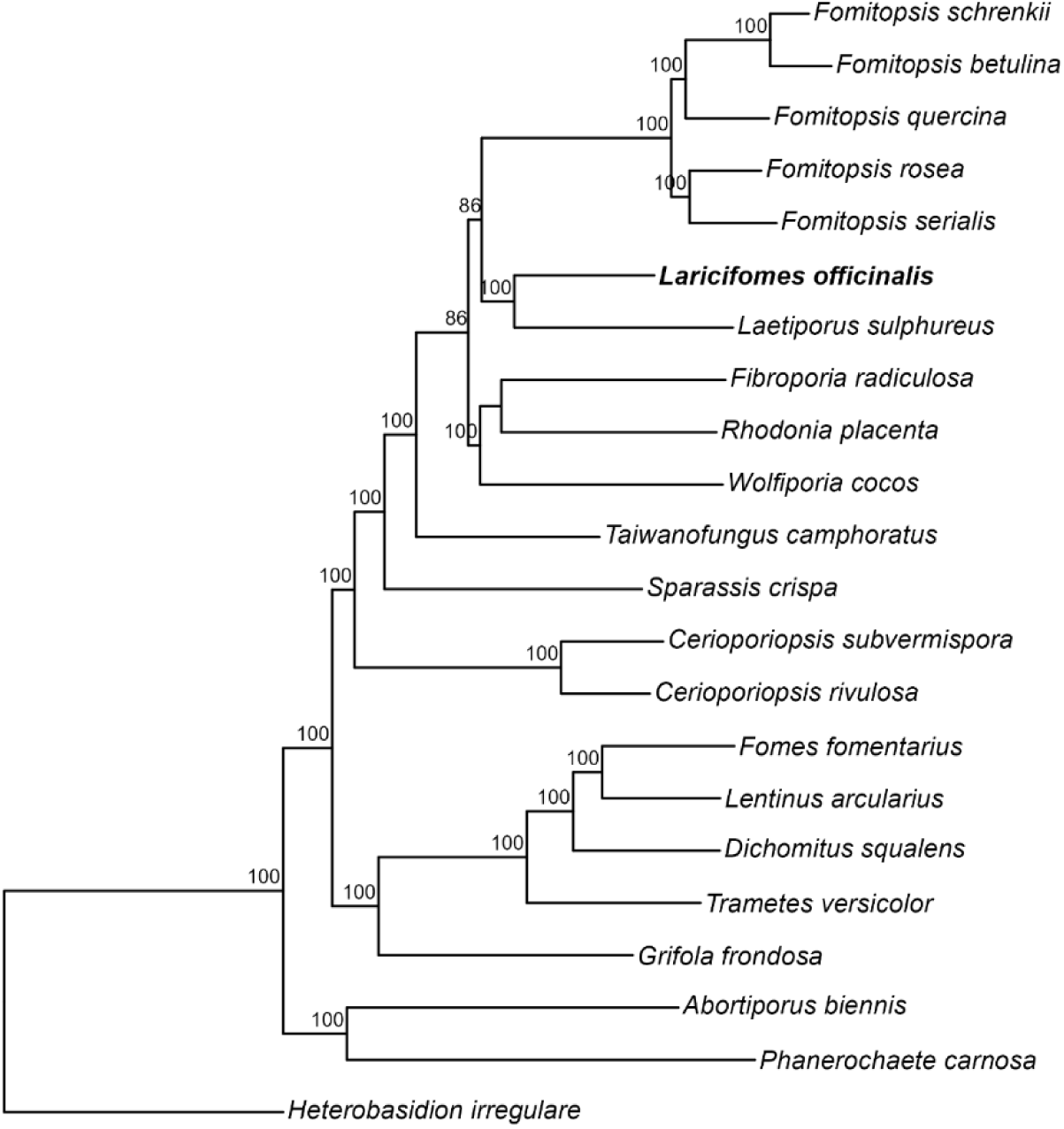
Maximum-likelihood phylogenetic tree constructed from a concatenated supermatrix alignment of amino acid sequences for 860 BUSCO genes shared among 22 taxa in the Polyporales. Branch support labels represent bootstrap values from 1000 ultra-fast bootstrap replicates.

After the phylogenomic analysis revealed a close taxonomic relationship between *L. officinalis* and *Lae. sulphureus*, collinearity among the largest scaffolds of each of their genomes was assessed using a synteny plot (Fig. 3).

**Fig. 3.**
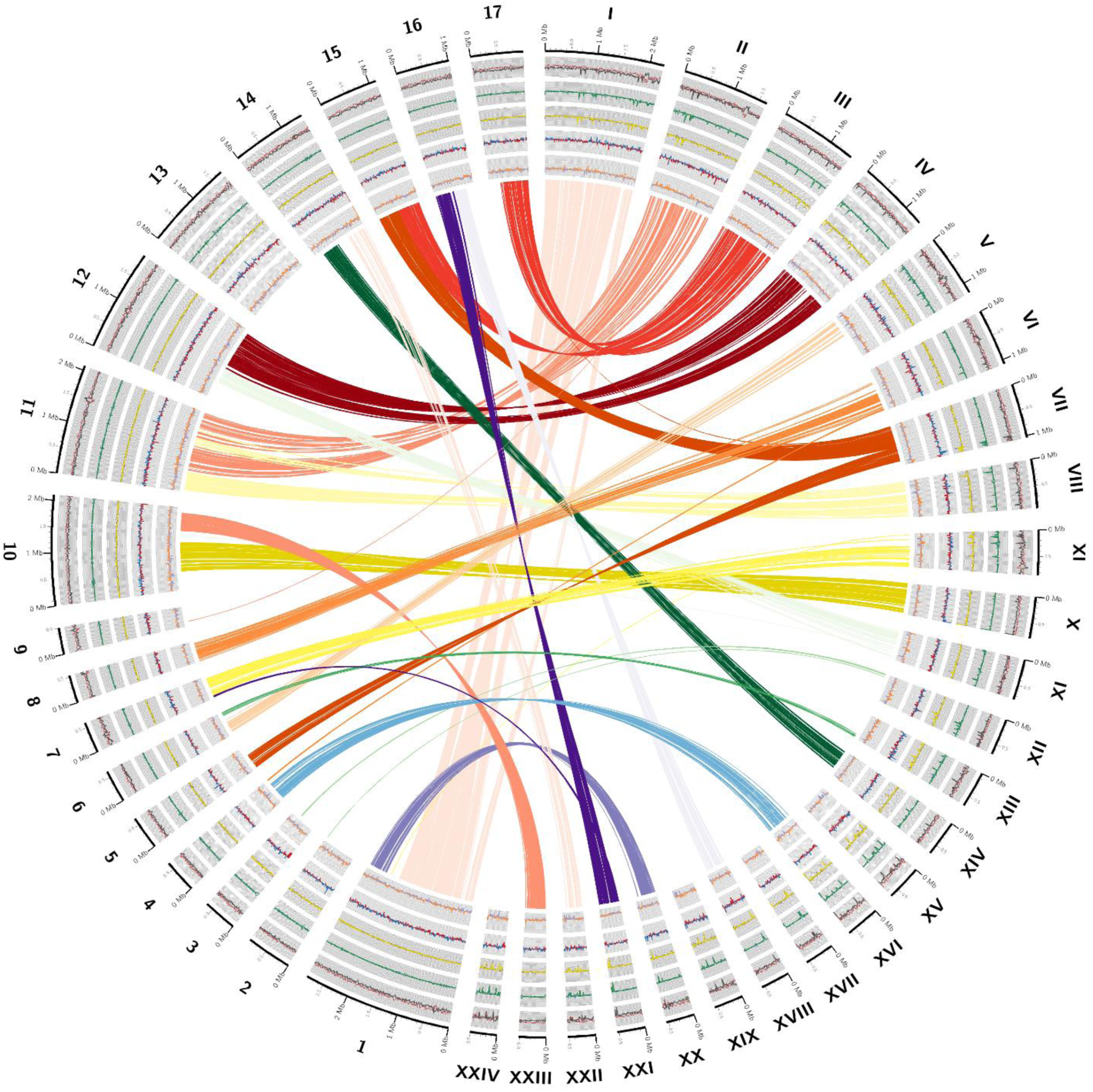
Synteny plot comparing the largest assembled genome scaffolds (>500 kb) from *Laricifomes officinalis* F01 (Arabic numerals) with those of *Laetiporus sulphureus* 93-53 (Roman numerals). Concentric subplots (from outer to inner) represent AT and GC nucleotide biases, GT and AC nucleotide biases, GA and CT nucleotide biases, GC skews, and AT skews. Colored ribbons indicate syntenic blocks shared among the scaffolds of each genome.

### Genome Size

The assembly showed high BUSCO completeness (99.4% and 93.7% for conserved basidiomycete genes and proteins, respectively), indicating a high-quality genome for a fungal lineage with limited characterization. Despite this, the agarikon genome is among the smallest reported for polypores in both assembly size and gene number. Of the 93 genome assemblies representing 81 species available in the Joint Genome Institute Polyporales dataset (MycoCosm) (Grigoriev et al. 2014), only *Fibroporia radiculosa* has a smaller assembly (28.4 Mb; 9,262 genes) (Tang et al. 2012). In comparison, *L. officinalis* encodes just 8,717 genes in 28.7 Mb, yielding a gene density that is generally consistent with other polypore fungi. While the evolutionary and ecological drivers and consequences of this reduced genome size remain unclear, possible explanations include lineage-specific loss of non-essential genes or streamlining associated with its specialization toward slow growth on long-lived old-growth conifer hosts.

### Mating Type Genes

Gene regions corresponding to the tetrapolar mating type loci, *matA* and *matB*, were identified on scaffolds 2 and 5 of the nuclear genome, respectively. The *matA* locus displayed homologs of many elements expected for this highly conserved region, including homeodomain-encoding (*HD*) mating type genes (AC3AMU_001439, AC3AMU_001440), a glycosyl transferase family 8 gene (*GLG2*) (AC3AMU_001435), a putative beta-flanking gene (*β-fg*) (AC3AMU_001438), and a mitochondrial intermediate peptidase gene (*MIP* = octapeptidyl aminopeptidase, *OCT1*) (AC3AMU_001441) (James et al. 2004; James et al. 2011; James et al. 2013; Dong et al. 2022). The *matB* locus comprises several pheromone mating factor G-protein coupled receptor (GPCR) genes (*STE3*) (AC3AMU_003118, AC3AMU_003122, AC3AMU_003123, AC3AMU_003124, AC3AMU_003125) and a major facilitator superfamily sugar transporter (*MFS1*) (AC3AMU_003119). Both mating loci were compared to those of *Lae. sulphureus* and *T. camphoratus* to evaluate synteny and collinearity across related species (Fig. 4). While there was broad collinearity among the genes present in the *matA* locus in *L. officinalis*, *T. camphoratus*, and *Lae. sulphureus*, the central HD genes did not meet the minimum alignment sequence identity (0.30), suggesting diversification among these genes, as would be expected due to their involvement in mating type specificity (James et al. 2011).

**Fig. 4.**
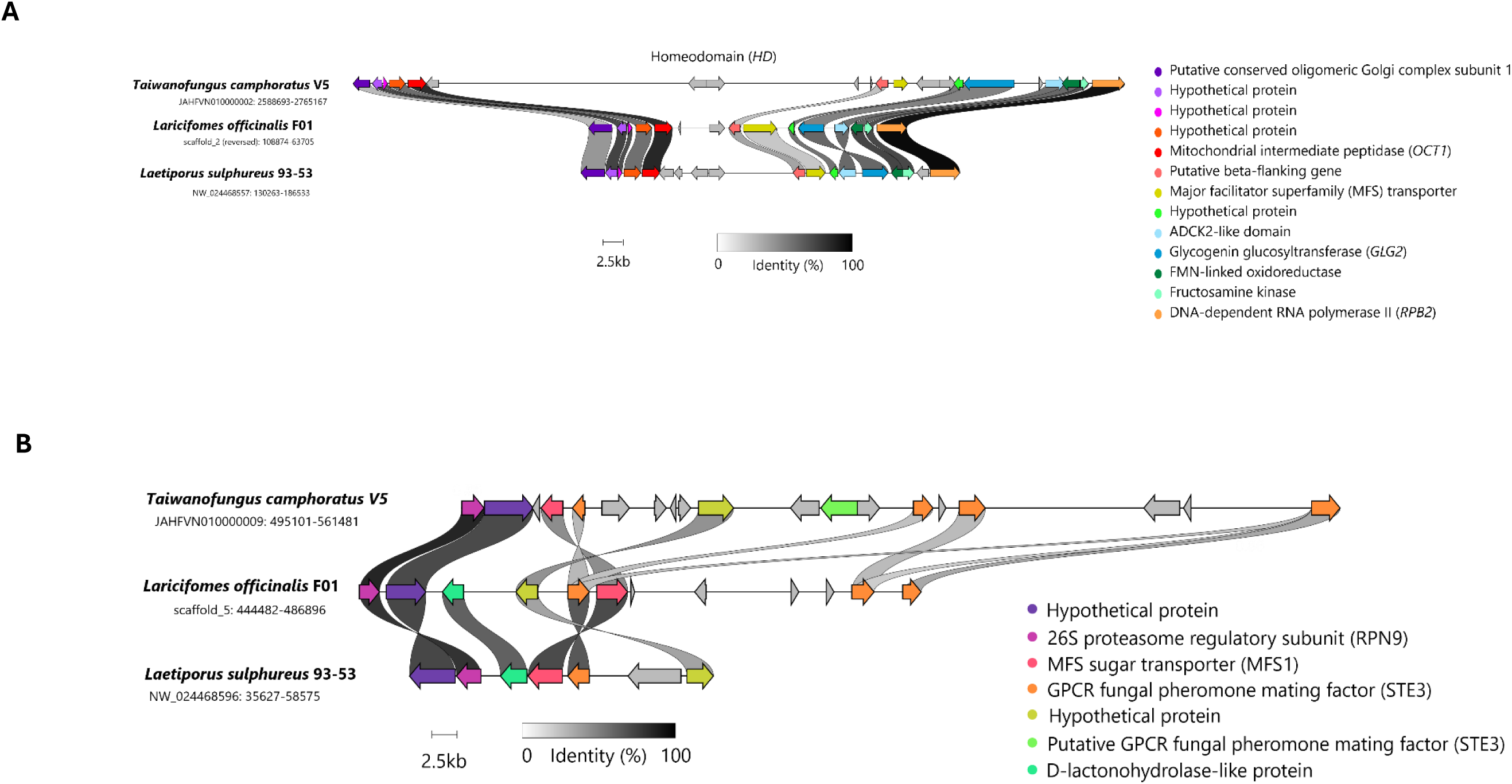
Synteny among gene clusters corresponding to the (a) *matA* and (b) *matB* loci from genome sequences of *Laricifomes officinalis* F01, *Laetiporus sulphureus* 93-53, and *Taiwanofungus camphoratus* V5.

### Carbohydrate Active Enzymes

A total of 223 genes encoding a suite of carbohydrate active enzymes (CAZymes) were identified in the nuclear genome of *L. officinalis*. This included 115 genes encoding glycoside hydrolases (GH), 52 encoding glycosyl transferases (GT), 39 associated with auxiliary activities (AA), 12 encoding carbohydrate esterases (CE), 3 encoding polysaccharide lyases (PL), and 2 encoding carbohydrate-binding modules (CBM) (Table 2). The numbers of CAZymes observed in each of these classes are generally consistent with other brown rot fungi such as *T. camphoratus*, which exhibits a similar ecological life history (Chen et al. 2022).

### Terpenoid Biosynthesis

The agarikon genome encodes numerous enzymes related to the biosynthesis of various terpenoids, which are commonly attributed for the bioactivities and medicinal benefits of polypore fungi (Lin et al. 2014; Saba et al. 2015; Kuo et al. 2016; Lin et al. 2017; Chen et al. 2022; Dong et al. 2022; Khalilov et al. 2022; Wang et al. 2023; Karunarathna et al. 2025). In particular, lanostane triterpenoids such as fomitopsins, officimalonic acids, sulphurenic acid, and eburicoic acid have been associated with medicinal effects including anti-tumor activity (Wu et al. 2004; Wu et al. 2009; Zhang et al. 2014; J. Han et al. 2016; Zhang et al. 2024; Zhang et al. 2025). Our analyses revealed several biosynthetic gene clusters (BGCs) involved in the mevalonate pathway and downstream triterpenoid synthesis, crucial to the production of ergosterol as well as secondary metabolites.

Specifically, we identified a BGC containing a farnesyl pyrophosphate (FPP) synthase (*ERG20*, AC3AMU_002446) and two BGCs encoding enzymes responsible for the dimerization of FPP into squalene, a bifunctional FPP farnesyltransferase/squalene synthase (*ERG9*, AC3AMU_005648) and a putative squalene synthase gene (AC3AMU_002051). Although the epoxidase (*ERG1*, AC3AMU_003390) required to convert squalene into oxidosqualene was found outside of a BGC, we also characterized a distinct BGC anchored on the lanosterol synthase gene (*ERG7*, AC3AMU_007627) (Fig. 5), which encodes an oxidosqualene cyclase responsible for converting oxidosqualene into lanosterol, an essential precursor for the production of diverse lanostane triterpenoids (Lin et al. 2015; Gressler et al. 2021; Zhang et al. 2025).

**Fig. 5.**
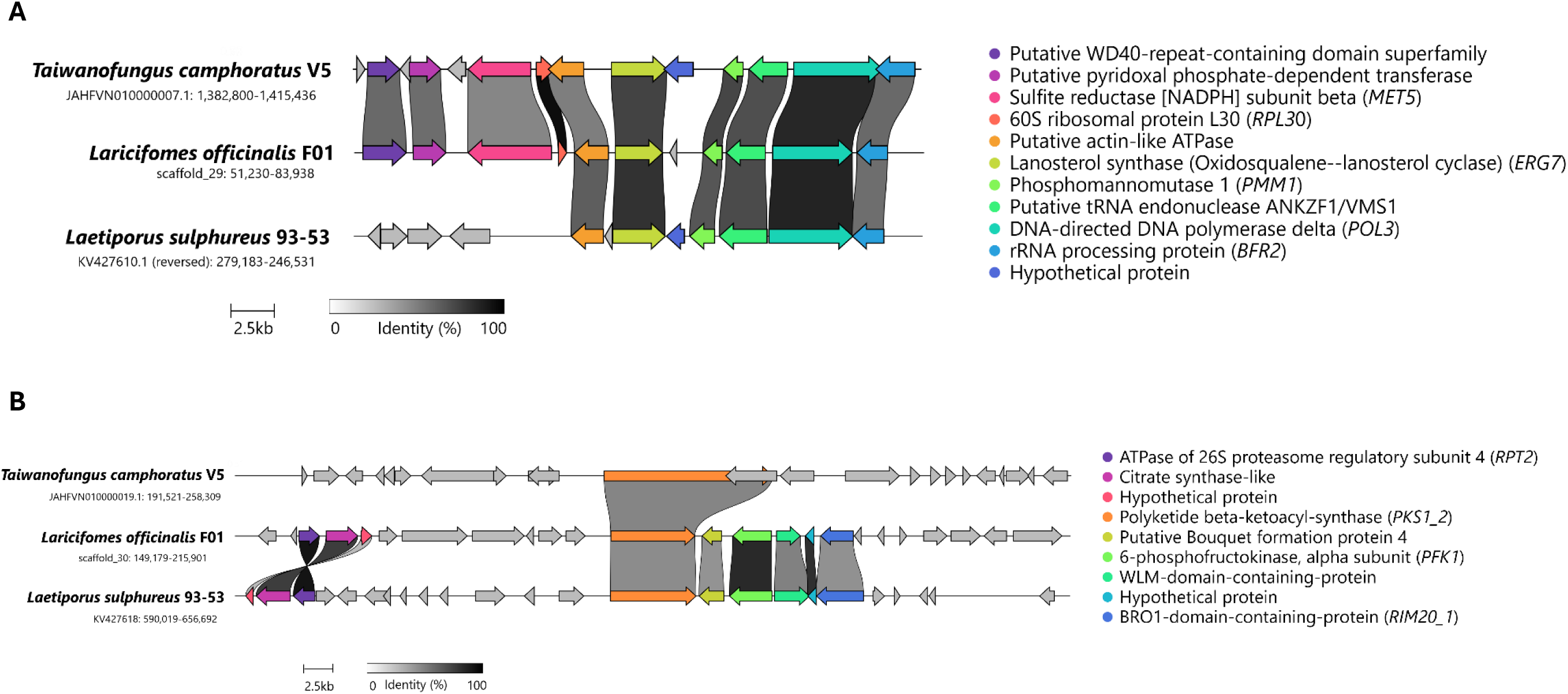
Synteny among secondary metabolite genes in biosynthetic gene clusters (BGCs) from the genomes of *Laricifomes officinalis* F01, *Laetiporus sulphureus* 93-53, and *Taiwanofungus camphoratus* V5. a) Genes encoding enzymes involved in the lanostane triterpenoid pathways, anchored on lanosterol synthase (*ERG7*). b) Genes in the BGC encoding a non-reducing PKS (NR-PKS) representing a putative homolog of an orsellinic acid biosynthesis cluster, anchored on the PKS gene.

We also identified a delta(24)-sterol C-methyltransferase (*ERG6,* AC3AMU_008380), a key enzyme involved in the production of ergosterol and certain lanostane triterpenoids (Dong et al. 2022; He et al. 2025; Zhang et al. 2025). The predicted amino acid sequence shows significant homology to a steroid methyltransferase (SMT) enzyme identified from metabolomic and transcriptomic analyses of *T. camphoratus* and may be functionally equivalent, with one proposed function as a catalyst of dehydrosulphurenic acid biosynthesis (Zhang et al., 2025). Others suggest this enzyme may catalyze the production of a key intermediary in eburicoic acid biosynthesis by *Lae. sulphureus* (Dong et al. 2022).

### Sesquiterpenes

Numerous additional terpene-related BGCs were identified, often containing multiple biosynthetic genes alongside cytochrome P450s (CYP450s) and transporters, suggesting a rich genomic repertoire for the biosynthesis of complex and diverse terpenoids. Among these, three BGCs contained genes encoding sesquiterpene synthases (*STS02,* AC3AMU_007975; *STS03,* AC3AMU_003641; *AGR3,* AC3AMU_005810) similar to those associated with the production of bicyclic sesquiterpenes including (+)-δ-cadinol by *Coniophora puteana* (Ringel et al. 2022), as well as α-muurolene, δ-cadinene, and γ-muurolene by *Cyclocybe* (= *Agrocybe*) *aegerita* (Zhang et al. 2020; Orban et al. 2021; Wang et al. 2023). These enzymes are structurally similar to Basidiomycete STS Clade I, which generally catalyze the production of α-muurolene and cadinols via C1,10 cyclization of FPP (Ichinose and Kitaoka 2018; Nagamine et al. 2019; Gressler et al. 2021; Wu et al. 2022; Wang et al. 2023).

Another terpene biosynthesis cluster contained two distinct *STS10* genes encoding sesquiterpene synthases (AC3AMU_003569 and AC3AMU_003571) that exhibited conserved amino acid motifs (DEXSD; NDVYSYNME) and Pfam annotations (PF19086) suggesting similarity to Clade II STS enzymes (Gressler et al. 2021; Wu et al. 2022; Wang et al. 2023). These enzymes, such as PpSTS10 from *R. placenta* and AcTPS4 from *T. camphoratus*, generally catalyze the production of cadinenes via C1,10 cyclization of nerolidyl pyrophosphate (NPP) (Ichinose and Kitaoka 2018; Nagamine et al. 2019; Gressler et al. 2021; Wu et al. 2022; Wang et al. 2023).

Two BGCs contained genes encoding Clade III STS enzymes homologous to those in *R. placenta*, identified based on conserved amino acid motifs. One was *STS14* (AC3AMU_005447), encoding a putative homolog of PpSTS14 with shared motifs DEYTD and NDIASYNKE, while the other was *STS09* (AC3AMU_007273), encoding an enzyme homologous to PpSTS09 with shared motifs DEYSD and NDMLSXNVE (Ichinose and Kitaoka 2018; Wang et al. 2023). Clade III STS enzymes catalyze a C1,11 cyclization of FPP to produce complex tricyclic backbones such as pentalenene and Δ^6^-protoilludene (Ichinose and Kitaoka 2018; Gressler et al. 2021; Wu et al. 2022; Wang et al. 2023). Collectively, STS enzymes in this clade are among the most important from a pharmaceutical perspective, as they generally catalyze the production of a wide array of secondary metabolites with potential antitumor, antibiotic, and anti-inflammatory activities (Gressler et al. 2021).

### Trichodiene Synthase Homologs

Another terpene BGC contained a terpene synthase gene (AC3AMU_001097) and two distinct copies of the monoterpene synthase gene *STS25* (AC3AMU_001102 and AC3AMU_001003), while a separate BGC similarly contained a homologous gene (AC3AMU_002333). Despite minor variation in these motifs among the four copies present in the *L. officinalis* F01 genome, each gene featured trichodiene synthase (TRI5) domains based on Pfam (PF06330) and InterPro (IPR024652) domains, as well as the presence of conserved amino acid motifs, including the Mg^2+^ binding site (DDXXX), a pyrophosphate and ion chelation region (NDXXSFYKEEL), and an FPP-binding site (XXRYRL) (Zhang et al. 2024). Trichodiene synthase catalyzes the cyclization of FPP to the bicyclic sesquiterpene trichodiene (Wang et al. 2023), but divergent evolution has yielded TRI5-like enzymes with potential to biosynthesize other terpenes (Cong et al. 2024). Zhang et al. (2024) identified six of these *Tri5*-like genes in the genome of *Taiwanofungus gaoligongensis* and demonstrated their homology to proteins in *Sparassis crispa* and *Trametes versicolor*. These enzymes may represent trichodiene synthase-like sesquiterpene synthases (TDTSs), which can catalyze the production of diverse monoterpenes and sesquiterpenes (Cong et al. 2024).

Although the functions of these *L. officinalis* STS25 enzymes have not been characterized, they are likely homologous to STS25 enzymes described from *R. placenta* (e.g., PpSTS25) and are similar to enzymes such as Fompi1, originally described from a North American isolate of *F. pinicola*, which was later determined to be *F. mounceae* (Wawrzyn et al. 2012; Haight et al. 2019; Spirin et al. 2024). Similarly, all eight predicted monoterpene synthases encoded in the *Lae. sulphureus* genome were homologous to *STS25* in *R. placenta* (Dong et al., 2022). These Clade IV STS enzymes catalyze a C1,6 cyclization of NPP to produce compounds such as α-cuprenene (Fompi1), as well as monoterpenes such as myrcene, and linalool (PpSTS25) (Ichinose and Kitaoka 2018; Gressler et al. 2021; Wu et al. 2022; Wang et al. 2023; Cong et al. 2024).

Additional BGCs identified by antiSMASH included two associated with the production of terpene precursors, each encoding putative FPP synthase-like polyprenyl synthetases (AC3AMU_005238 and AC3AMU_006708). Another BGC involved in terpene biosynthesis included a haloacid-dehalogenase-like hydrolase gene clustered with two CYP450 monooxygenase genes and a short-chain dehydrogenase/reductase (SDR). Collectively, these results suggest that *L. officinalis* has the potential to produce a structurally diverse collection of terpenoid compounds.

### Polyketide Biosynthesis

One large region near the edge of scaffold_28 was predicted by antiSMASH to contain a type 1 polyketide synthase (T1PKS) BGC amid three neighboring NRPS-like clusters, although it is unclear whether the apparent overlap between these reflects a functionally integrated biosynthetic region or low-confidence boundary prediction. This T1PKS cluster centered on a highly-reducing beta-ketoacyl synthase (*pks1_1,* AC3AMU_007588) includes associated genes predicted to encode an AMP-dependent acyl ligase (AC3AMU_007594), a crotonyl-CoA reductase (AC3AMU_007593), a flavin-dependent halogenase (AC3AMU_007590), a Hotdog-fold thioesterase (AC3AMU_007585), multiple CYP450s (AC3AMU_007582, AC3AMU_007587, AC3AMU_007589), multiple MFS1 proton-linked monocarboxylate transporters (AC3AMU_007580, AC3AMU_007581, AC3AMU_007586, AC3AMU_007595), and an intermembrane transporter (AC3AMU_007591). Additional genes in the adjacent predicted NRPS-like clusters similarly encode AMP-dependent synthetase/ligases, additional CYP450s, and MFS1 transporters, as well as a D-lactate ferricytochrome c oxidoreductase (*DLD1_3*, AC3AMU_007579), an acyl-CoA dehydrogenase/oxidase (AC3AMU_007578), and a GMC oxidoreductase (AC3AMU_007572).

The presence of a putative flavin-dependent halogenase (AC3AMU_007590) in this T1PKS cluster raises the possibility that this region might contribute to the biosynthesis of chlorinated coumarins, which were previously discovered in the same *L. officinalis* isolate (F01) during an investigation of agarikon’s antibiotic activity against *Mycobacterium tuberculosis* (Hwang et al., 2013). However, while the halogenase features a tryptophan fold commonly associated with aromatic substrate halogenation, the PKS (AC3AMU_007588) encodes a typical highly reducing PKS (HR-PKS) with a KS-AT-DH-KR-(NAD) domain architecture, which generally indicates aliphatic (rather than aromatic) polyketide synthesis. Consistent with most fungal HR-PKSs (Chooi and Tang 2012), this PKS also lacks a C-terminal reductase or thioesterase domain, indicating that polyketide chain release and downstream processing are likely mediated by trans-acting enzymes encoded in the region. In addition to the adjacent Hotdog-fold thioesterase (AC3AMU_007585) which could mediate the primary chain release, a pair of functionally equivalent adenylation enzymes with A-PP-TD domain architecture (AC3AMU_007594, AC3AMU_007605) could provide an alternative or subsequent route for additional post-PKS tailoring of metabolites produced by this BGC.

Together, these trans-acting enzymes provide a plausible mechanism for the decoupling of initial polyketide chain assembly and final backbone formation, potentially explaining how a single HR-PKS could support the production of multiple secondary metabolites with diverse structures, including aromatic compounds. Furthermore, oxidative transformations by the numerous CYP450s and redox enzymes in the region could also mediate aromatic cyclization of following aliphatic polyketide synthesis, analogous to noncanonical fungal biosynthetic mechanisms in which reduced polyketides are converted to aromatic products through extensive post-PKS oxidative remodeling (Liu et al. 2019). Although the enzymes encoded within this T1PKS BGC are consistent with the formation, oxidative tailoring, potential halogenation, and export of polyketides, the specific enzymatic steps and metabolic product(s) remain speculative; furthermore, any potential functional involvement of the adjacent NRPS-like clusters remains to be determined.

### Non-Reducing Polyketide Synthases

A second T1PKS BGC identified by antiSMASH contains a beta-ketoacyl synthase (*pks1_2*, AC3AMU_007745), encoding a non-reducing PKS (NR-PKS) with the canonical domain structure SAT-KS-AT-PT-CP-CP-TE (Gressler et al. 2021). A homology search with BLASTp, along with synteny analysis via clinker, suggests this PKS is a putative homolog of PKS63787, an orsellinic acid synthase confirmed to be involved in the biosynthesis of antroquinonol meroterpenoids, benzenoids, and benzoquinones in *T. camphoratus* (Yu et al. 2016; Chou et al. 2017; Gressler et al. 2021; Kuang et al. 2021; Zhang et al. 2024). In particular, *T. camphoratus* antroquinonols have been studied extensively, with substantial preclinical evidence demonstrating their anti-cancer and anti-inflammatory effects achieved through multiple pathways (Kuang et al. 2021), including by inhibiting nitric oxide, demonstrated following treatment of LPS-induced macrophages with multiple antroquinonols derived from the mycelium (Yang et al. 2009).

Homologs of this NR-PKS cluster were previously identified from the genomes of *T. gaoligongensis*, *T. camphoratus*, *Lae. sulphureus*, *Fomitopsis palustris*, and *F. quercina* (Zhang et al. 2024). The NR-PKS cluster identified in the genome of *L. officinalis* F01 is structurally similar to those of *Lae. sulphureus*, *F. palustris*, and *F. quercina*, which, unlike *Taiwanofungus* spp., all contain genes encoding modifying proteins including a 6-phosphofructokinase (AC3AMU_007747), a WLM-domain-containing-protein (AC3AMU_007748), and BRO1-domain-containing-proteins (*RIM20_1*, FUN007750 and *RIM20_2*, FUN007751) in addition to the core PKS gene (Zhang et al. 2024). However, in the *L. officinalis* genome, this cluster lacks the CYP450 genes that are present in the homologous clusters in *Lae. sulphureus* (Fig. 5b) and *F. palustris* (Zhang et al. 2024).

In addition to the terpene synthases, HR-PKS, and NR-PKS clusters described above, other terpene synthases and NRPS-like clusters were also identified by antiSMASH. These warrant further investigation as to their identity and homology with other fungal secondary metabolite clusters identified previously. Despite the fact that *L. officinalis* has a rather compact genome by comparison to other fungi in the Polyporales, the relative number and diversity of SM BGCs encoded in its genome is comparable to those found in the often much larger genomes of other polypores, though with fewer PKS clusters than are generally observed (Chen et al. 2022; Dong et al. 2022; Zhang et al. 2024). While some of the identified SM BGCs have similarities to those described from other fungi, others appear to be potentially novel variants that may allow *L. officinalis* to produce a unique suite of secondary metabolites to which agarikon’s numerous demonstrated medicinal benefits may be attributable.

## CONCLUSION

Here we present highly contiguous and extensively annotated nuclear and mitochondrial genomes of agarikon (*Laricifomes officinalis*), an endangered fungus that is the sole member of a unique evolutionary lineage. In addition to resolving the core mitochondrion genome, potentially one of the largest among polypores, annotation of the relatively small 28.8 Mbp nuclear genome revealed 8,717 total genes, including 8,604 protein-coding genes. A genome-wide BUSCO phylogeny clarified agarikon’s taxonomic position within the Polyporales, confirming it does not form a monophyletic group with members of the genus *Fomitopsis*. While comparative genomic analyses revealed its affinity to *Laetiporus sulphureus*, it also showed similarities to *Taiwanofungus camphoratus* despite its more distant evolutionary relationship. Characteristics of both mating loci (*matA* and *matB*) were consistent with the tetrapolar mating system expected for a polypore fungus. Despite having one of the smallest nuclear genomes among fungi in the Polyporales, *L. officinalis* encodes 310 genes in 27 biosynthetic gene clusters with the potential to produce a diverse array of terpenoids, polyketides, and other bioactive compounds. These could represent an untapped reservoir of potentially novel metabolites that may have medicinal value. Creating a robust reference genome for *L. officinalis* is a foundational step for understanding the biology of this unique fungus, which will enable future population genomics studies and facilitate functional characterization of secondary metabolites underlying the medicinal benefits of agarikon that have been reported for centuries.

## ACKNOWLEDGEMENTS

We gratefully acknowledge Jim Gouin, David Sumerlin, Travis Zalesky, and Jacqueline Morgado, whose contributions to the collection and preservation of agarikon isolates were integral to the completion of this work.

## DATA AVAILABILITY

All genomic data and associated metadata have been deposited in NCBI GenBank under BioProject PRJNA1460884 and BioSample SAMN57657796. Sequence data have been deposited in the Short Read Archive (SRA) with accession numbers SRR38352409 (ONT long reads) and SRR38352408 (Illumina short reads). The annotated mitochondrial have been deposited in GenBank with accession numbers PZ428821 (mitochondrial genome sequence). The nuclear genome has been submitted to GenBank with submission number SUB16160987 and is currently being processed by NCBI. All data will become publicly available coinciding with publication of the peer-reviewed manuscript.

All software code for reproducing the analyses and figures presented herein will be made publicly available in the following GitHub repository: https://github.com/Patrick-Bennett/Agarikon-Genomics.

## SUPPLEMENTARY MATERIALS

Supplementary Figure 1. Clusters of orthologous groups (COGs) from the annotated Laricifomes officinalis F01 genome.

Supplementary Figure 2. Top 30 most abundant Pfam domains from the annotated Laricifomes officinalis F01 genome.

Supplementary Figure 3. Top 30 gene ontology (GO) terms in each category from the annotated *Laricifomes officinalis* F01 genome.

Supplementary Table 1. Sequence data QC statistics for Illumina reads from the SeqKit analysis.

Supplementary Table 2. Sequence data QC statistics for Oxford Nanopore reads from the SeqKit analysis.

Supplementary Table 3. Quality assessment of nuclear genome assembly from QUAST following polishing with Illumina reads via Pypolca.

Supplementary Table 4. Repeat content predicted from the nuclear genome of Laricifomes officinalis F01.

Supplementary Table 5. Summary of mitochondrial genome annotations for Laricifomes officinalis F01.

Supplementary Table 6. Polyporales genomes used in the phylogenomic and/or comparative analyses with the nuclear genome *Laricifomes officinalis* F01.

