## Supplementary Figures and Tables for "Genomic Characterization of the Endangered Medicinal Polypore Agarikon (*Laricifomes officinalis* syn. *Fomitopsis officinalis*)"

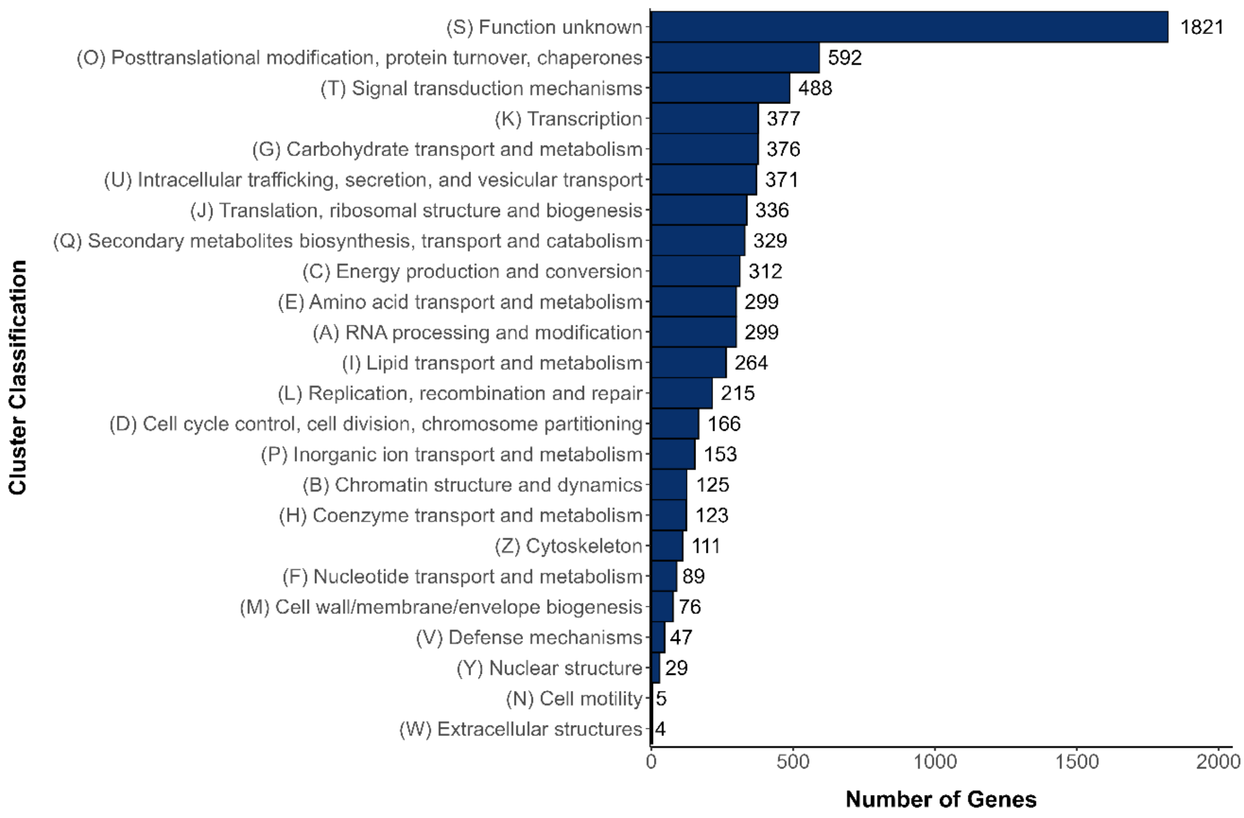


**Suppl. Fig. 1.** Clusters of orthologous groups (COGs) from the annotated nuclear genome of *Laricifomes officinalis* F01.

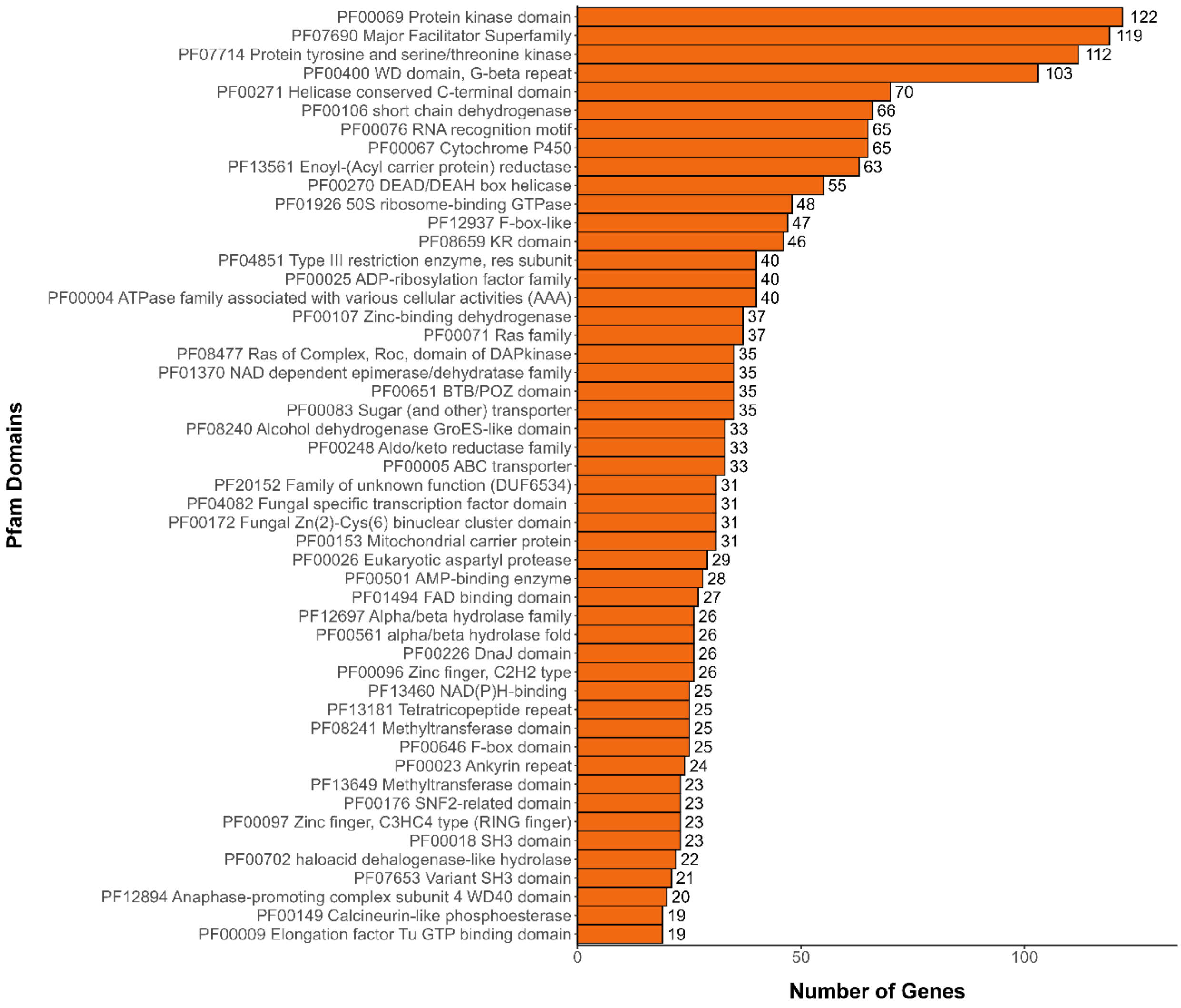


**Suppl. Fig. 2.** Top 30 most abundant Pfam domains from the annotated nuclear genome of *Laricifomes officinalis* F01.

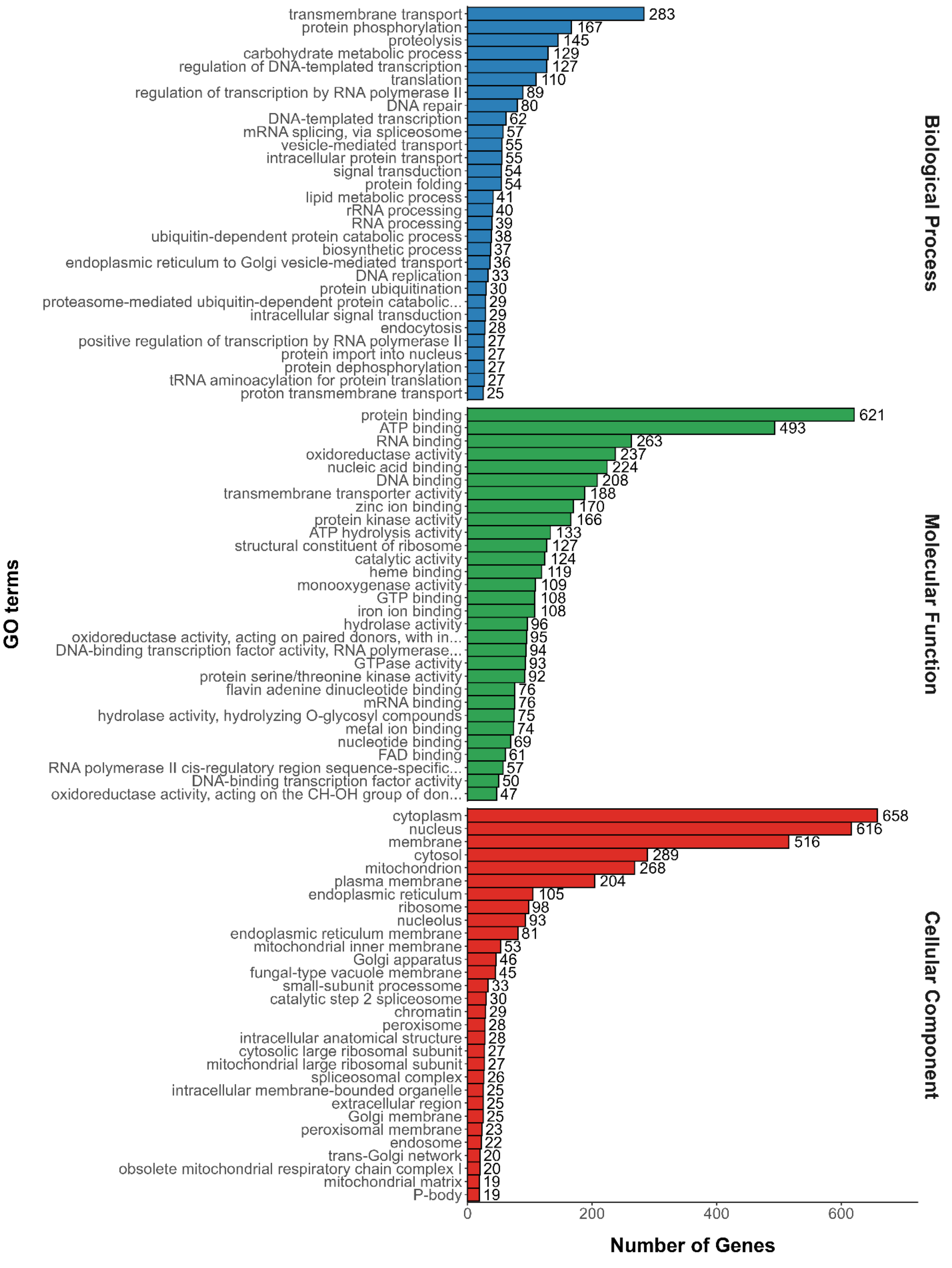


**Suppl. Fig. 3.** Top 30 gene ontology (GO) terms in each category from the annotated nuclear genome of *Laricifomes officinalis* F01.

**Supplementary Table 1.** Sequence data QC statistics for Illumina reads from the SeqKit analysis.

| File | num_seqs[a] | sum_len[b] | min_len[c] | avg_len[d] | max_len[e] | Q1[f] | Q2[g] | Q3[h] | sum_gap[i] | N50[j] | N50_num[k] | Q20(%)[l] | Q30(%)[m] | AvgQual[n] | GC(%)[o] | sum_n[p] |
|---|---|---|---|---|---|---|---|---|---|---|---|---|---|---|---|---|
| R1 | 11,646,036 | 1,746,905,400 | 150 | 150 | 150 | 150 | 150 | 150 | 0 | 150 | 1 | 97.69 | 92.83 | 26.62 | 43.54 | 11,144 |
| R2 | 11,646,036 | 1,746,905,400 | 150 | 150 | 150 | 150 | 150 | 150 | 0 | 150 | 1 | 96.34 | 89.77 | 24.82 | 43.69 | 7,129 |

**Supplementary Table 2.** Sequence data QC statistics for Oxford Nanopore reads from the SeqKit analysis.

| num_seqs | sum_len | min_len | avg_len | max_len | Q1 | Q2 | Q3 | sum_gap | N50 | N50_num | Q20(%) | Q30(%) | AvgQual | GC(%) | sum_n |
|---|---|---|---|---|---|---|---|---|---|---|---|---|---|---|---|
| 115,538 | 554,888,901 | 147 | 4,802.7 | 65,783 | 1,559 | 3,420 | 6,882 | 0 | 7,726 | 9,511 | 41.73 | 10.12 | 9.8 | 47.47 | 0 |

[a] number of sequences
[b] number of bases or residues, with gaps or spaces counted
[c] minimal sequence length, with gaps or spaces counted
[d] average sequence length, with gaps or spaces counted
[e] maximum sequence length, with gaps or spaces counted
[f] first quartile of sequence length, with gaps or spaces counted
[g] median of sequence length, with gaps or spaces counted
[h] third quartile of sequence length, with gaps or spaces counted
[i] number of gaps
[j] N50 length
[k] L50 (N50 number)
[l] percentage of bases with quality score greater than 20
[m] percentage of bases with quality score greater than 30
[n] average quality
[o] percentage of GC content
[p] number of ambiguous letters (N, n, X, x)

**Supplementary Table 3.** Quality assessment of nuclear genome assembly from QUAST following polishing with Illumina reads via Pypolca.

| | |
|---|---|
| # contigs | 66 |
| # contigs (>= 0 bp) | 66 |
| # contigs (>= 1000 bp) | 66 |
| # contigs (>= 5000 bp) | 65 |
| # contigs (>= 10000 bp) | 64 |
| # contigs (>= 25000 bp) | 61 |
| # contigs (>= 50000 bp) | 53 |
| Largest contig | 2838896 |
| Total length | 28757392 |
| Total length (>= 0 bp) | 28757392 |
| Total length (>= 1000 bp) | 28757392 |
| Total length (>= 5000 bp) | 28752609 |
| Total length (>= 10000 bp) | 28742679 |
| Total length (>= 25000 bp) | 28685031 |
| Total length (>= 50000 bp) | 28411918 |
| N50 | 997481 |
| N90 | 204582 |
| auN | 1200039 |
| L50 | 9 |
| L90 | 34 |
| GC (%) | 51.96 |
| # N's per 100 kbp | 0 |
| # N's | 0 |

**Supplementary Table 4.** Repeat content predicted from the nuclear genome of *Laricifomes officinalis* F01.

| Type | Number of elements | Length occupied | Percentage of sequence |
|---|---|---|---|
| Retroelements | 923 | 2275524 bp | 7.91% |
| SINEs: | 0 | 0 bp | 0.00% |
| Penelope | 72 | 144265 bp | 0.50% |
| LINEs: | 72 | 144265 bp | 0.50% |
| CRE/SLACS | 0 | 0 bp | 0.00% |
| L2/CR1/Rex | 0 | 0 bp | 0.00% |
| R1/LOA/Jockey | 0 | 0 bp | 0.00% |
| R2/R4/NeSL | 0 | 0 bp | 0.00% |
| RTE/Bov-B | 0 | 0 bp | 0.00% |
| L1/CIN4 | 0 | 0 bp | 0.00% |
| LTR elements: | 851 | 2131259 bp | 7.41% |
| BEL/Pao | 0 | 0 bp | 0.00% |
| Ty1/Copia | 82 | 202102 bp | 0.70% |
| Gypsy/DIRS1 | 735 | 1886424 bp | 6.56% |
| Retroviral | 0 | 0 bp | 0.00% |
| DNA transposons | 103 | 49038 bp | 0.17% |
| hobo-Activator | 0 | 0 bp | 0.00% |
| Tc1-IS630-Pogo | 56 | 21600 bp | 0.08% |
| En-Spm | 0 | 0 bp | 0.00% |
| MuDR-IS905 | 0 | 0 bp | 0.00% |
| PiggyBac | 0 | 0 bp | 0.00% |
| Tourist/Harbinger | 0 | 0 bp | 0.00% |
| Other (Mirage, P-element, Transib) | 0 | 0 bp | 0.00% |
| Rolling-circles | 0 | 0 bp | 0.00% |
| Unclassified: | 1989 | 573550 bp | 1.99% |
| Total interspersed repeats: | | 2898112 bp | 10.08% |
| Small RNA: | 4 | 68554 bp | 0.24% |
| Satellites: | 0 | 0 bp | 0.00% |
| Simple repeats: | 0 | 0 bp | 0.00% |
| Low complexity: | 0 | 0 bp | 0.00% |
| Total bases masked: | | 2966666 bp | 10.32 % |

* most repeats fragmented by insertions or deletions have been counted as one element

**Supplementary Table 5.** Summary of mitochondrial genome annotations for *Laricifomes officinalis* F01.

| Gene | Product | Start | End | Introns | Intron Type(s) |
|---|---|---|---|---|---|
| **ATP Synthase Genes** | | | | | |
| atp6_1 | ATP synthase F0 subunit a | 137868 | 138113 | 0 | |
| atp6_2 | ATP synthase F0 subunit a | 159151 | 158372 | 0 | |
| atp8 | ATP synthase F0 subunit 8 | 167643 | 167485 | 0 | |
| atp9 | ATP synthase F0 subunit c | 144142 | 143921 | 0 | |
| **Cytochrome B and Cytochrome C Oxidase Genes** | | | | | |
| cob | apocytochrome b | 175395 | 194964 | 11 | Group I (IA, IB, ID, derived) |
| cox1 | cytochrome c oxidase subunit 1 | 41 | 35289 | 23 | Group I (IA, IB, ID, derived); Group II |
| cox2 | cytochrome c oxidase subunit 2 | 85104 | 96670 | 6 | Group I (IB, IC1, ID, derived) |
| cox3 | cytochrome c oxidase subunit 3 | 100711 | 104194 | 2 | Group I (IB, derived) |
| **NADH Dehydrogenase Genes** | | | | | |
| nad1 | NADH dehydrogenase subunit 1 | 168651 | 173779 | 4 | Group I (derived) |
| nad2 | NADH dehydrogenase subunit 2 | 144365 | 148179 | 1 | Group I (IB) |
| nad3 | NADH dehydrogenase subunit 3 | 148195 | 148590 | 0 | |
| nad4 | NADH dehydrogenase subunit 4 | 38215 | 39726 | 0 | |
| nad4L | NADH dehydrogenase subunit 4L | 104784 | 105263 | 0 | |
| nad5 | NADH dehydrogenase subunit 5 | 106348 | 117773 | 7 | Group I (IB, ID, IC2, derived) |
| nad6 | NADH dehydrogenase subunit 6 | 125488 | 126204 | 0 | |
| **Mitochondrial rRNA Gene** | | | | | |
| rps3 | ribosomal protein S3 | 119740 | 120747 | 0 | |
| rnl | large subunit rRNA | ~ 55159 (+/- 50 nt) | uncertain | 1 | Group I (IC2) |
| rns | small subunit rRNA | ~ 76036 | ~ 83366 | ~ 4 | Group I (IC2, IB); Group II |
| **Mitochondrial tRNA Genes** | | | | | |
| trnA(tgc) | tRNA Alanine | 44221 | 44291 | | |
| trnC(gca) | tRNA Cysteine | 168023 | 168098 | | |
| trnD(gtc) | tRNA Aspartate | 167204 | 167132 | | |
| trnE(ttc) | tRNA Glutamate | 197273 | 197343 | | |
| trnF(gaa) | tRNA Phenylalanine | 44475 | 44546 | | |
| trnG(tcc) | tRNA Glycine | 99277 | 99348 | | |

| | | | |
|---|---|---|---|
| trnH(gtg) | tRNA Histidine | 153251 | 153323 |
| trnI(gat) | tRNA Isoleucine | 175078 | 175148 |
| trnK(ttt) | tRNA Lysine | 45160 | 45231 |
| trnL(taa) | tRNA Leucine | 196892 | 196976 |
| trnL(tag) | tRNA Leucine | 83984 | 84066 |
| trnM(cat) | tRNA Methionine | 46421 | 46494 |
| trnM(cat) | tRNA Methionine | 150031 | 150103 |
| trnM(cat) | tRNA Methionine | 197051 | 197122 |
| trnN(gtt) | tRNA Asparagine | 196660 | 196732 |
| trnP(tgg) | tRNA Proline | 196457 | 196530 |
| trnQ(ttg) | tRNA Glutamine | 44789 | 44862 |
| trnR(cct) | tRNA Arginine | 142318 | 142248 |
| trnR(tcg) | tRNA Arginine | 84273 | 84344 |
| trnR(tct) | tRNA Arginine | 128477 | 128549 |
| trnR(tct) | tRNA Arginine | 168258 | 168328 |
| trnS(gct) | tRNA Serine | 159672 | 159592 |
| trnS(tga) | tRNA Serine | 46783 | 46868 |
| trnT(tgt) | tRNA Threonine | 44698 | 44769 |
| trnV(tac) | tRNA Valine | 83417 | 83487 |
| trnW(cca) | tRNA Tryptophan | 167354 | 167427 |
| trnY(gta) | tRNA Tyrosine | 45867 | 45950 |
| **Open Reading Frames (ORFs)** | | | |
| orf673 | hypothetical protein | 134 | 2155 |
| orf809 | hypothetical protein | 2567 | 4996 |
| orf345 | GIY | 5139 | 6176 |
| orf342 | LAGLIDADG | 6542 | 7570 |
| orf317 | LAGLIDADG | 8039 | 8992 |
| orf180 | LAGLIDADG | 9378 | 9920 |
| orf156 | LAGLIDADG | 11624 | 12094 |
| orf380 | LAGLIDADG | 12189 | 13331 |
| orf166 | LAGLIDADG | 13516 | 14016 |
| orf113 | hypothetical protein | 15290 | 15631 |

| | | | |
|---|---|---|---|
| orf170 | LAGLIDADG | 16135 | 16647 |
| orf127 | hypothetical protein | 17065 | 17448 |
| orf359 | LAGLIDADG | 17976 | 19055 |
| orf319 | LAGLIDADG | 19297 | 20256 |
| orf311 | LAGLIDADG | 20865 | 21800 |
| orf322 | LAGLIDADG | 21870 | 22838 |
| orf413 | LAGLIDADG | 23009 | 24250 |
| orf186 | LAGLIDADG | 26764 | 27324 |
| orf263 | LAGLIDADG | 27380 | 28171 |
| orf734 | hypothetical protein | 28613 | 30817 |
| orf234 | LAGLIDADG | 31132 | 31836 |
| orf173 | GIY | 32290 | 32811 |
| orf312 | GIY | 33780 | 34718 |
| orf54 | LAGLIDADG | 39947 | 40111 |
| orf42 | LAGLIDADG | 40129 | 40257 |
| orf176 | hypothetical protein | 49674 | 50204 |
| orf135 | hypothetical protein | 50441 | 50848 |
| orf219_1 | hypothetical protein | 51557 | 50898 |
| orf419 | hypothetical protein | 53211 | 51952 |
| orf131_1 | hypothetical protein | 53828 | 53433 |
| orf276 | hypothetical protein | 54830 | 54000 |
| orf285 | LAGLIDADG | 57013 | 57870 |
| orf77 | LAGLIDADG | 57874 | 58107 |
| orf60 | LAGLIDADG | 58101 | 58283 |
| orf303 | LAGLIDADG | 59112 | 60023 |
| orf351 | LAGLIDADG | 60871 | 61926 |
| orf434 | LAGLIDADG | 62529 | 63833 |
| orf207 | LAGLIDADG | 64824 | 65447 |
| orf325 | LAGLIDADG | 66190 | 67167 |
| orf250 | LAGLIDADG | 68850 | 69602 |
| orf214 | GIY | 71008 | 71652 |
| orf100 | hypothetical protein | 72046 | 72348 |

| | | | |
|---|---|---|---|
| orf103 | GIY | 72299 | 72610 |
| orf45 | LAGLIDADG | 76957 | 77094 |
| orf88 | LAGLIDADG | 77098 | 77364 |
| orf76 | LAGLIDADG | 77593 | 77823 |
| orf252 | LAGLIDADG | 78969 | 79727 |
| orf164 | hypothetical protein | 80187 | 80681 |
| orf231 | LAGLIDADG | 80626 | 81321 |
| orf161_1 | hypothetical protein | 81566 | 82051 |
| orf243 | LAGLIDADG | 81948 | 82679 |
| orf77 | LAGLIDADG | 82692 | 82925 |
| orf166_2 | hypothetical protein | 85338 | 85838 |
| orf336 | hypothetical protein | 85906 | 86916 |
| orf219 | LAGLIDADG | 87482 | 88141 |
| orf238 | hypothetical protein | 91413 | 92129 |
| orf452 | GIY | 95158 | 96516 |
| orf269 | GIY | 97061 | 97870 |
| orf323 | LAGLIDADG | 100930 | 101901 |
| orf223 | hypothetical protein | 103434 | 104105 |
| orf124_1 | hypothetical protein | 106739 | 107113 |
| orf172 | LAGLIDADG | 107176 | 107694 |
| orf232 | hypothetical protein | 108422 | 109120 |
| orf261 | hypothetical protein | 109164 | 109949 |
| orf183 | LAGLIDADG | 110646 | 111197 |
| orf168 | LAGLIDADG | 111403 | 111909 |
| orf251 | hypothetical protein | 112130 | 112885 |
| orf199 | hypothetical protein | 113391 | 113990 |
| orf156_2 | hypothetical protein | 121766 | 121296 |
| orf110 | hypothetical protein | 122981 | 122649 |
| orf245 | hypothetical protein | 123727 | 122990 |
| orf163 | LAGLIDADG | 126192 | 126683 |
| orf452_2 | hypothetical protein | 128895 | 130253 |
| orf147_1 | hypothetical protein | 131531 | 131088 |

| | | | |
|---|---|---|---|
| orf479 | hypothetical protein | 132342 | 133781 |
| orf430 | hypothetical protein | 134305 | 135597 |
| orf173_2 | hypothetical protein | 137327 | 137848 |
| orf147_2 | hypothetical protein | 138819 | 139262 |
| orf105 | hypothetical protein | 140380 | 140063 |
| orf140 | hypothetical protein | 141963 | 141541 |
| orf194 | GIY | 143939 | 143355 |
| orf124_2 | hypothetical protein | 145807 | 146181 |
| orf190 | hypothetical protein | 146619 | 147191 |
| orf102 | hypothetical protein | 151342 | 151034 |
| orf638 | hypothetical protein | 155417 | 153501 |
| orf129 | hypothetical protein | 162272 | 161883 |
| orf220 | hypothetical protein | 162744 | 163406 |
| orf161_2 | hypothetical protein | 163705 | 164190 |
| orf474 | hypothetical protein | 166127 | 164703 |
| orf327 | LAGLIDADG | 170370 | 171353 |
| orf133 | hypothetical protein | 171374 | 171775 |
| orf75 | GIY | 173967 | 174194 |
| orf107 | LAGLIDADG | 177596 | 177919 |
| orf169 | hypothetical protein | 177982 | 178491 |
| orf449 | GIY | 179067 | 180416 |
| orf131 | GIY | 182273 | 182668 |
| orf513 | LAGLIDADG | 184065 | 185606 |
| orf280 | LAGLIDADG | 186029 | 186871 |
| orf123 | LAGLIDADG | 190552 | 190923 |
| orf101 | hypothetical protein | 194040 | 194345 |

**Supplementary Table 6.** Polyporales genomes used in the phylogenomic and/or comparative analyses with the nuclear genome of *Laricifomes officinalis* F01.

| Taxon | GenBank Accession | Citation |
|---|---|---|
| *Abortiporus biennis* | GCA_022606235.1 | (Hage et al. 2021) |
| *Dichomitus squalens* | GCF_000275845.1 | (Floudas et al. 2012) |
| *Fibroporia radiculosa* | GCF_000313525.1 | (Tang et al. 2012) |
| *Fomes fomentarius* | GCA_022606135.1 | (Hage et al. 2021) |
| *Fomitopsis betulina* | GCA_022606075.1 | (Hage et al. 2021) |
| *Fomitopsis quercina* | GCA_001632345.1 | (Nagy et al. 2016) |
| *Fomitopsis schrenkii* | GCA_000344655.2 | (Floudas et al. 2012) |
| *Fomitopsis serialis* | GCF_022376445.1 | (Hage et al. 2021) |
| *Cerioporiopsis subvermispora* | GCA_000320605.2 | (Fernandez-Fueyo et al. 2012) |
| *Grifola frondosa* | GCA_001683735.1 | (Min et al. 2016) |
| *Heterobasidion irregulare* | GCF_000320585.1 | (Olson et al. 2012) |
| *Laetiporus sulphureus* | GCF_001632365.1 | (Nagy et al. 2016) |
| *Cerioporiopsis rivulosa* | GCA_001687445.1 | (Miettinen et al. 2016) |
| *Phanerochaete carnosa* | GCF_000300595.1 | (Suzuki et al. 2012) |
| *Lentinus arcularius* | GCA_004369055.1 | (Varga et al. 2019) |
| *Rhodofomes roseus* | GCF_022264815.1 | (Hage et al. 2021) |
| *Rhodonia placenta* | GCF_002117355.1 | (Gaskell et al. 2017) |
| *Sparassis crispa* | GCF_003851025.1 | (Kiyama et al. 2018) |
| *Taiwanofungus camphoratus* | GCA_022598655.1 | (Chen et al. 2022) |
| *Trametes versicolor* | GCF_000271585.1 | (Floudas et al. 2012) |
| *Wolfiporia cocos* | GCA_000344635.1 | (Floudas et al. 2012) |

Chen C-L et al. 2022. Sexual crossing, chromosome-level genome sequences, and comparative genomic analyses for the medicinal mushroom *Taiwanofungus camphoratus* (Syn. *Antrodia cinnamomea*, *Antrodia camphorata*). Microbiology Spectrum. 10(1):e02032-21. https://doi.org/10.1128/spectrum.02032-21

Fernandez-Fueyo E et al. 2012. Comparative genomics of *Ceriporiopsis subvermispora* and *Phanerochaete chrysosporium* provide insight into selective ligninolysis. Proceedings of the National Academy of Sciences. 109(14):5458–5463. https://doi.org/10.1073/pnas.1119912109

Floudas D et al. 2012. The Paleozoic origin of enzymatic lignin decomposition reconstructed from 31 fungal genomes. Science. 336(6089):1715–1719. https://doi.org/10.1126/science.1221748

Gaskell J et al. 2017. Draft genome sequence of a monokaryotic model brown-rot fungus *Postia* (*Rhodonia*) *placenta* SB12. Genomics Data. 14:21–23. https://doi.org/10.1016/j.gdata.2017.08.003

Hage H et al. 2021. Gene family expansions and transcriptome signatures uncover fungal adaptations to wood decay. Environmental Microbiology. 23(10):5716–5732. https://doi.org/10.1111/1462-2920.15423

Kiyama R, Furutani Y, Kawaguchi K, Nakanishi T. 2018. Genome sequence of the cauliflower mushroom *Sparassis crispa* (Hanabiratake) and its association with beneficial usage. Sci Rep. 8(1):16053. https://doi.org/10.1038/s41598-018-34415-6

Miettinen O et al. 2016. Draft Genome Sequence of the White-Rot Fungus *Obba rivulosa* 3A-2. Genome Announcements. 4(5):10.1128/genomea.00976-16. https://doi.org/10.1128/genomea.00976-16

Min B et al. 2016. Whole genome sequencing of *Grifola frondosa* 9006-11. https://www.ncbi.nlm.nih.gov/datasets/genome/GCA_001683735.1/

Nagy LG et al. 2016. Comparative genomics of early-diverging mushroom-forming fungi provides insights into the origins of lignocellulose decay capabilities. Mol Biol Evol. 33(4):959–970. https://doi.org/10.1093/molbev/msv337

Olson Å et al. 2012. Insight into trade-off between wood decay and parasitism from the genome of a fungal forest pathogen. New Phytologist. 194(4):1001–1013. https://doi.org/10.1111/j.1469-8137.2012.04128.x

Suzuki H et al. 2012. Comparative genomics of the white-rot fungi, *Phanerochaete carnosa* and *P. chrysosporium*, to elucidate the genetic basis of the distinct wood types they colonize. BMC Genomics. 13(1):444. https://doi.org/10.1186/1471-2164-13-444

Tang JD et al. 2012. Short-read sequencing for genomic analysis of the brown rot fungus *Fibroporia radiculosa*. Applied and Environmental Microbiology. 78(7):2272–2281. https://doi.org/10.1128/AEM.06745-11

Varga T et al. 2019. Megaphylogeny resolves global patterns of mushroom evolution. Nat Ecol Evol. 3(4):668–678. https://doi.org/10.1038/s41559-019-0834-1
